# Contrasting Rhizosphere Nitrogen Dynamics in Andropogoneae Grasses: Implications for Sustainable Agriculture

**DOI:** 10.1101/2024.06.03.597142

**Authors:** Sheng-Kai Hsu, Bryan D Emmett, Alexandria Haafke, Germano Costa-Neto, Aimee J Schulz, Nicholas Lepak, Thuy La, Taylor AuBuchon-Elder, Charles Hale, Sierra S Raglin, Jonathan O Ojeda-Rivera, Angela D Kent, Elizabeth A Kellogg, M Cinta Romay, Edward S Buckler

**Affiliations:** Institute for Genomic Diversity, Cornell University, Ithaca, NY, USA 14853; USDA-ARS, National Laboratory for Agriculture and the Environment, Ames, IA, USA 50011; Section of Plant Breeding and Genetics, Cornell University, Ithaca, NY USA 14853; USDA-ARS, Robert W. Holley Center for Agriculture and Health, Ithaca, NY, USA 14853; Donald Danforth Plant Science Center, St. Louis, MO, USA 63132; Carl R. Woese Institute for Genomic Biology, University of Illinois at Urbana-Champaign, Urbana, IL, USA 61801; Department of Natural Resources and Environmental Sciences, University of Illinois at Urbana-Champaign, Urbana, IL, USA 61801; Arnold Arboretum of Harvard University, Boston, MA, USA 02130

**Author notes:** Correspondence: Sheng-Kai Hsu,; Edward S Buckler.

**Keywords:** Rhizosphere, Nitrogen Cycle, Andropogoneae, Sustainability, Evolution

## Abstract

**Background:** Nitrogen (N) fertilization in crop production significantly impacts ecosystems, often disrupting natural plant-microbe-soil interactions and causing environmental pollution. Our research tested the hypothesis that phylogenetically related perennial grasses might preserve rhizosphere management strategies conducive to a sustainable N economy for crops.

**Method:** We analyzed the N cycle in the rhizospheres of 36 Andropogoneae grass species related to maize and sorghum, investigating their impacts on N availability and losses. This assay is supplemented with the collection and comparison of native habitat environment data for ecological inference as well as cross-species genomic and transcriptomic association analyses for candidate gene discovery.

**Result:** Contrary to our hypothesis, all examined annual species, including sorghum and maize, functioned as N "Conservationists," reducing soil nitrification potential and conserving N. In contrast, some perennial species enhanced nitrification and leaching ("Leachers"). Yet a few other species exhibited similar nitrification stimulation effects but limited NO_3_^-^ losses ("Nitrate Keepers"). We identified significant soil characteristics as influential factors in the eco-evolutionary dynamics of plant rhizospheres, and highlighted the crucial roles of a few transporter genes in soil N management and utilization.

**Conclusion:** These findings serve as valuable guidelines for future breeding efforts for global sustainability.

## Introduction

The development of inorganic nitrogen (N) fertilizer has revolutionized modern agriculture (Khush, 2001; Smith *et al*., 2020). Nevertheless, the benefit comes at the cost of severe environmental impacts. The soil microbial processes of nitrification and denitrification, which transform the less mobile ammonium (NH_4_ ^+^) into free nitrate (NO_3_^-^) and nitrite (NO_3_^-^), lead to inefficient use of synthesized inorganic N on farms and environmental pollution (Bremner & Blackmer, 1978; Schlesinger, 2009; Billen *et al*., 2013). NO_3_^-^ leaches from agricultural soil, leading to eutrophication of neighboring water systems (Schlesinger, 2009; Billen *et al*., 2013). The hypoxic zone in the Gulf of Mexico serves as a cautionary example (Rabalais & Turner, 2019). Soil nitrous oxide (N_2_O and NO_x_) emission accounts for ∼60% of agricultural greenhouse gas footprint: N_2_O exhibits 265 times higher global warming potential than CO_2_, and 75% of anthropogenic N_2_O emission in the US is from nitrification and denitrification processes in agricultural soils (EPA, 2023). N loss from agricultural soils is particularly evident in early spring (Lu *et al*., 2022), likely due to the mismatch in timing between crop fertilization, plant N demand, and soil microbial activities (Supplementary Figure S1; Bender *et al*., 2013; Hartman *et al*., 2022; Kosola *et al*., 2023). Therefore, ensuring early, efficient and conservative utilization of soil N on farms is crucial for global sustainability.

Recently, Subbarao & Searchinger, 2021 proposed a “more ammonium solution,” which emphasizes managing soil nitrification rates to conserve N as NH_4_^+^ in the soil. The proposed approach requires careful and precise orchestration of the three-way interactions among plants, soil and microbes. Soil characteristics, such as moisture and pH, affect plant growth, microbial activity, and their interactions, including competition and various symbiotic relationships (McNear, 2013). Conversely, microbes and plants can also modify soil properties to benefit their own growth (McNear, 2013). Given the complexities of plant-soil-microbe interactions, it is not entirely clear whether and how to control nitrification in agricultural soil.

Previous efforts have focused on plants and their impact on rhizosphere microbes. Crop scientists identified the biological nitrification inhibition (BNI) capacity in various crops (Subbarao *et al*., 2007c, 2009, 2013; Pariasca Tanaka *et al*., 2010; Tesfamariam *et al*., 2014; Sun *et al*., 2016; Byrnes *et al*., 2017; Nuñez *et al*., 2018; Villegas *et al*., 2020), with the goal of improving their ability to inhibit soil nitrification on farms. Significant advances were made in identifying root exudates with nitrification-inhibiting effects and understanding their biosynthesis (Tesfamariam *et al*., 2014; Widhalm & Rhodes, 2016; Wang *et al*., 2020; Pan *et al*., 2021; Otaka *et al*., 2022). More recently, a few studies have started to investigate the genetic variation within crop species to explore the potential of genetic improvement (Petroli *et al*., 2023) and between species to seek transferable BNI-contributing alleles from wild relatives (Subbarao *et al*., 2007b, 2021).

However, the benefit of BNI capacity in a crop species could vary depending on its NH_4_ ^+^ uptake efficacy (Abalos *et al*., 2018) and its tolerance to NH_4_ ^+^ toxicity (Esteban *et al*., 2016). There is massive variation in the preference between NH_4_ ^+^ and NO_3_^-^ among plant species adapted to distinct environments (Houlton *et al*., 2007; Kahmen *et al*., 2008; Boudsocq *et al*., 2012; Britto & Kronzucker, 2013). Hence, it is conceivable that suppression of nitrification can be detrimental to NO_3_^-^-preferring species (Boudsocq *et al*., 2012; Konaré *et al*., 2019) and the nitrification process is not necessarily harmful to the environment if the converted NO_3_^-^ can be efficiently assimilated and utilized. For instance, even within species, two ecotypes with opposing effects on nitrification were found in different sites for *Hyparrhenia diplandra* (Lata *et al*., 2000). We hypothesize that diverse species adapting independently to various environments manage their rhizosphere differently, resulting in divergent patterns of N-cycling. Amongst the range in rhizosphere N dynamics across species, we seek alternative strategies for N conservation in agricultural soils. In particular, we anticipate finding favorable phenotypes among perennial grasses that establish earlier in the season than the annual crops.

In this study, we focus on the Andropogoneae tribe of grasses, which has evolved to dominate 17% of global land area with C4 photosynthesis, adapting to diverse habitats, often forming large populations (Moore *et al*., 2019; Cowan *et al*., 2020; Bachle *et al*., 2022). Importantly, maize, one of the most productive crops on earth, belongs to this tribe. We carried out a systematic assay on the rhizosphere N traits in 36 Andropogoneae species, including maize and sorghum, during their active vegetative growing stage. We discovered pronounced phenotypic variation and identified three distinct rhizosphere N management strategies amongst different species: NH_4_^+^ conservation, NO_3_^-^ leaching, and NO_3_^-^ retention. Further biogeographical and environmental association provides insights into the ecological selection forces shaping the three strategies. Through cross-species genomic/transcriptomic association analysis, we identified key genes and breeding targets that could enhance agricultural sustainability.

## Materials and Methods Plant materials

In this study we investigate a total of 36 Andropogoneae species, including two important annual crops: maize (*Zea mays*) and sorghum (*Sorghum bicolor*) (Supplementary Table S1). These 36 species are distributed across the Andropogoneae phylogeny (Figure 1a). Significant BNI effect was shown in maize and sorghum previously (Subbarao *et al*., 2013; Otaka *et al*., 2022; Petroli *et al*., 2023). Except for maize and sorghum (seeds produced in the lab nursery), live clones of the other 34 species were either clonally propagated from wild grassland habitats or grown from seeds obtained from wild collections, the USDA-ARS National Plant Germplasm System, Iowa State University, or commercial sources (Supplementary Table S1). Plant materials were clonally propagated in a greenhouse with 14 hours of daylight at 28°C and 10 hours of nighttime at 22°C. Supplemental lights turned on when natural light was below 500 W/m^2^. Soil moisture was maintained by daily watering. The plants were fertilized three times a week during watering (480ppm 21-5-20: 7.92% NH_4_ ^+^ and 13.08% NO_3_^-^; 1350ppm 15-5-15 + 4% Ca + 2% Mg: 3% NH_4_^+^ and 12% NO_3_^-^; 300ppm Fe chelate).

**Figure 1.**
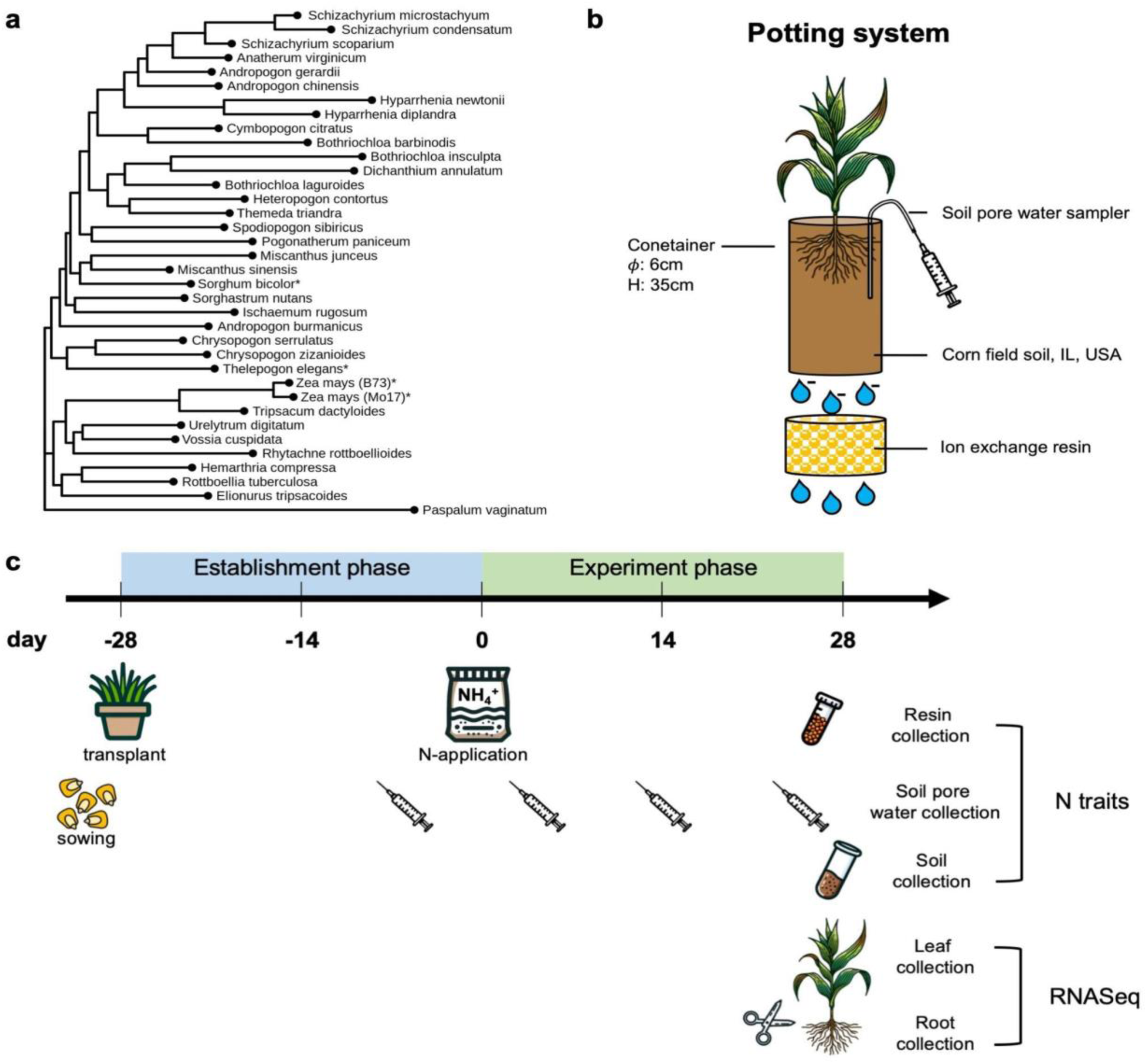
Plant materials and experimental design. a. Phylogenetic tree of the Andropogoneae grass species studied in the study. Asterisks denote annual taxa. **b.** Illustration of the potting setup. Each seedling is planted in a tall cylindrical pot with a height (H) of 35cm and a diameter (𝟇) of 6cm. A soil pore water sampler is inserted to the soil and a trap with ion-exchange resins is installed beneath the pot. **c.** Timeline of the greenhouse experiment. There are four weeks of establishment phase for the plant to recover from the transplantation. The experiment starts with the application of 150 kg/ha NH_4_^+^-N as (NH_4_)_2_SO_4_ (1000 ppm N). We collect soil pore water samples on day -4, 3, 15, 24 of the experiment. On day 27, resin traps are collected. On day 28, we collected soil samples and plant tissues. Soil pore water, resin and soil samples are for rhizosphere N trait measurement and the plant tissues are subjected to RNA extraction and sequencing.

## Greenhouse experiment

We performed a greenhouse experiment to investigate the phylogenetic variation of the rhizosphere N cycle across Andropogoneae. This experiment followed a complete randomized design (CRD) with three replicates per species. Greenhouse condition was the same as the maintenance. For mesocosm soil, we used a 1:4 mixture of US corn belt soil (sourced from IL, USA, 40.05180278°N 88.23133889°W; Soil features: OM 3.0%, CEC 16.4 meq/100g, pH 6.9, NO_3_^-^ 5.9 ppm,NH_4_^+^ 2.5 ppm, P 22 ppm, K 109 ppm) and Cornell Potting mix (a premix of 3.8 ft^2^ peat, 4 ft^2^ vermiculite, 4 ft^2^ perlite, 50 lbs turface, 3 lbs limestone, 4 lbs 10-5-10 Media Mix fertilizer, and 2 lbs calcium sulfate). Maize and sorghum were germinated and grown for two weeks. Then, seedlings of maize and sorghum and clonal plantlets of other species were transplanted to cylindrical containers (diameter of 6 cm and height of 35 cm) filled with 700g of mesocosm soil (Figure 1b). The clonal plantlets were trimmed to 15cm above soil to promote regrowth. After transplantation, we allowed a 4-week establishment period with regular nutrient supply as in the maintenance protocol. Afterward, a one-time application of 150 kg/ha NH_4_^+^-N as (NH_4_)_2_SO_4_ (1000 ppm N) initiates the formal experiment (Figure 1c). At the end of the establishment period, fine nylon mesh bags containing 400ml of ion exchange resins (MAG-MB, Resintech) are placed under the containers to capture leached NO_3_^-^ from the system, and Rhizon soil moisture samplers (Eijkelkamp Agrisearch Equipment) were installed for soil pore water sampling (Figure 1b). After transplantation, we irrigated each plant with 100ml of water daily to control for the leaching rate. We collected soil pore water samples on day -4, 3, 15, 24 of the experiment (Figure 1c). Sampling is conducted an hour after irrigation of 100ml water by suction pressure using a common medical syringe via the Rhizon samplers. A sample of 10 ml of soil solution was collected and stored at -20°C for later assay. On day 27, resin traps were collected (Figure 1c). The resin was thoroughly homogenized and an aliquot of 40 ml resin of each bag was sampled and stored at 4°C. On day 28, we collected root (5cm from the tips), leaf (5cm from the tips) and soil samples (Figure 1c). Two samples of 40ml sieved soil per replicate were collected and stored at 4°C: one for NO_3_^-^ extraction and one for potential nitrification rate measurement. Both collected soil and resin samples were processed within 14 days after the sampling.

## NO_3_^-^ extraction and quantification

We extracted 2 grams of fresh soil media in 5 ml of 2M KCl extraction solution. The samples were shaken for 1 hour. After shaking, the samples were allowed to rest undisturbed for at least 30 minutes. The top layer of the extract solution, in a volume greater than 1 ml, was then filtered into new tubes. Similarly, for resin extraction, we extracted 2 grams of resin in 5 ml of 2M KCl. The extracts were stored at -80°C until shipment and quantification. Nitrate concentrations in pore water, soil and resin extracts were quantified colorimetrically using the VCl_3_/Griess method ^(^Mirand^a^ *et al*., 2001) in a 96-well microplate (Doane & Horwáth, 2003). Plates were incubated overnight in the dark and absorbance at 540 nm read using a Synergy HT microplate reader. A standard curve of 0 to 15 ppm was included on each plate and calculated concentrations expressed on a soil dry weight basis (µg N g soil^-1^).

## Potential nitrification rate measurement

Nitrification potential of mesocosm soil was assessed on fresh soils within a week of sampling using the shaken slurry method of (Hart *et al*., 1994). Approximately 5 g of moist soil media was placed in a half pint mason jar along with 33 ml of buffer (0.3 mM KH_2_PO_4_, 0.7 mM K_2_HPO_4_, 0.75 mM (NH_4_)SO_4_, pH 7.2), capped with a vented lid and shaken at 150 rpm at 30 ℃ for 24 hr. At 2, 4, 22 and 24 hr a 1 ml aliquot was sampled from each jar and centrifuged at 16,000 x g at 4°C for 10 min, the supernatant removed and stored at -20°C until quantification of nitrate as described above. Potential nitrification rate (PNR) was calculated as the rate of nitrate accumulation over time using the equation:

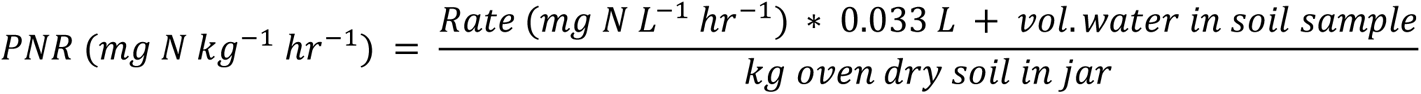

Comparing the PNR of each sample to bulk soil blank samples, we calculated the relative PNR in percentage as following:

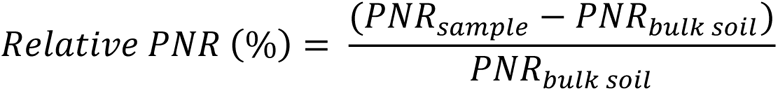

## Characterization of species with distinct rhizosphere N managing strategies

Comparing the average rhizosphere N trait measures of each species to the average measures of bulk soil blank samples, we identified three groups of species that manage their rhizosphere N differently. The groups and criteria are as following:

1. Conservationists: PNR reduced by 25% or more and NO_3_^-^ lost by <50%
2. Leachers: PNR elevated by 25% or more and NO_3_^-^ lost by >50%
3. Nitrate keepers: PNR elevated by 25% or more but NO_3_^-^ lost by >50%

## Characterization of geographic adaptation and environmental vectors (envPCs)

To estimate the natural environmental conditions in which each species occurs, we conducted a two-step environmental characterization. The initial step estimated the geographic range of each species using geographic coordinates sourced from species diversity and global distribution databases. This was accomplished using the R packages ‘BIEN’ v1.2.6 (Maitner, 2023) to access the Botanical Information and Ecology Network (BIEN). Then, the second step was focused on obtaining the environmental features characteristic of their adaptation. For each coordinate sample, we extracted a set of 94 environmental factors from diverse publications (Ross *et al*., 2018; Lembrechts *et al*., 2022) and databases, including WorldClim (Fick & Hijmans, 2017), FAO- GAEZ (‘GAEZ v4 Data Portal’) and GDSE (Shangguan *et al*., 2014) (Supplementary Table S2) using the packages ‘terra’ v1.7 (Hijmans *et al*., 2023). Subsequently, we computed the ranges of these environmental features for each species in terms of quantiles (10%, 50%, and 90%) across their geographic distribution. A total of 282 environmental features (a combination of environmental factors and quantiles) were obtained (Supplementary Table S3). Data quality control was performed by removing features with a rate of missing values higher than 10%, and imputing missing values using the function imputePCA() from the package ’missMDA’ v1.19 (Husson & Josse, 2023). Finally, using the function PCA() from the package ‘FactoMineR’ v2.9 (Husson *et al*., 2023), we did an eigen decomposition of the environmental relationship matrix to obtain linear and orthogonal combinations of numerous environmental features, the so-called environmental principal components (envPCs; Supplementary Table S3). We assumed that each envPC captures a different spatial and climatic trend of the global environmental diversity, serving as a proxy for the ecological range and the selection pressure that might shape the local-adaptation of each species.

## Association between habitat environment and rhizosphere N traits

To investigate the potential ecological selection pressure that shapes the distinct rhizosphere N dynamics in Andropogoneae species, we modeled the impact of different environmental features of the native habitat range on the rhizosphere N traits (i.e. potential nitrification rate, soil NO_3_^-^ content, and NO_3_^-^ loss) as follows:

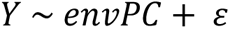

Where Y stands for average trait values, LoF denotes the binary scores that predict potential functional loss of the homolog, and 𝜀 is the residual.

Using a linear regression approach, we identified the envPCs that significantly co-vary with at least one rhizosphere N trait (p-value < 0.05). For each significant envPC, we extracted the significantly contributing environmental features and investigated the distribution of these variables in the three species groups.

## Genome assemblies

25 of the studied taxa have high quality long read assemblies available publicly (*Zea mays ssp. mays* var. B73 (Hufford *et al*., 2021), *Zea mays ssp. mays* var. Mo17 (Sun *et al*., 2018), *Sorghum bicolor* var. Btx623 (v3) (McCormick *et al*., 2018), *Miscanthus sinensis (Mitros et al., 2020)*, PanAnd (In prep.)). For 9 of the remaining species, short-read whole genome sequencing data were generated to supplement the long-read data. DNA was extracted from leaves. (Qiagen Inc., Germantown, MD). Extracted samples were quantified and Illumina Tru-Seq or nano Tru-seq libraries were constructed according to sample concentration. Samples were sequenced in pools of 24 individuals in one lane of an S4 flowcell in an Illumina Novaseq 6000 System with 150 bp pair-end reads. With short sequencing reads passing our customized quality control (Schulz *et al*., 2023), assemblies were generated using Megahit v1.2.9 (Li *et al*., 2015) using a minimum kmer size (--k-min) of 31 and default setting for the other parameters (Schulz *et al*., 2023). Two studied genomes without genome assemblies were excluded for phylogenetic and cross-species association analysis.

## Identification of the functional homolog

As the unit of the cross-species genomic and transcriptomic association analyses, we identified the most likely functional homologous genes among genome assemblies of various Andropogoneae taxa, using the closest outgroup, seashore paspalum (*Paspalum vaginatum*) (Sun *et al*., 2022) as reference. Annotated proteins in seashore paspalum were queried against all Andropgoneae genomes using miniProt v0.10 (Li, 2023). We retained the primary alignment (with the least sequence divergence from reference) in each genome to represent the functional homologs. We studied the functional homologs instead of the syntenic orthologs to prevent the comparison among orthologs that underwent differential genome contraction processes following whole genome duplication and/or polyploidization events, which occur repeatedly within the Andropogoneae tribe (Estep *et al*., 2014; Mitros *et al*., 2020; Zhang *et al*., 2024). Ideally, the functional homologs represent the most conserved gene copies among the homeologs/paralogs for each species. Multiple sequence alignments (MSAs) of the coding sequences (CDS) of each functional homolog were generated using MAFFT v7.508 (Katoh *et al*., 2002).

## Phylogenetic tree construction

We used the diversity of the putatively neutral loci to construct the phylogenetic relationship amongst the assayed species in this study. We used the four-fold degenerate sites in the CDS MSA of 2,000 random homologs that are present in >75% of all assayed species to construct the maximum likelihood species tree using RAxML v8.2.12 (Stamatakis, 2014) with the GTR model + Gamma correction for rate heterogeneity (-m GTRGAMMA).

## Loss of function prediction

Assuming functional constraint, for a homolog, we inferred that taxa with no alignment or unusually long genetic distance to the outgroup (one standard deviation away from average genetic distance) likely lost or shifted their function. Hence, we define a binary loss-of-function (LoF) score to represent the presence of a functional gene copy in each taxon. LoF score equals 0 for the taxa with missing or distant alignments while 1 for the other taxa.

## RNA extraction and sequencing

RNA was extracted from leaf or root tissues using the Direct-zol-96 RNA extraction kit by Zymo Research and RNeasy Plant mini kit by Qiagen based on manufacturers’ protocols. A few random RNA samples were selected for fragment analysis using the BioAnalyzer to check for fragment size and purity using RIN number of >6. RNA-seq library preparation was performed using the NEBNext® Ultra II Directional RNA Prep Kit for Illumina and NEBNext® Poly(A) mRNA Magnetic Isolation Module. Initial RNA concentration for each sample used ranged from 250- 1000ng. Pair-end 150bp reads were generated by the Illumina NovaSeq X Plus PE150 platform.

## Homolog expression quantification and normalization

Using Salmon v1.10.2 (Patro *et al*., 2017), we quantified the expression level of each homolog in different species in reference to the phylogenetically closest long-read genomes (Supplementary Table S1). In addition to coding exons of each homolog, we included reads mapped to putative untranslated regions (within 500bp up- and downstream of the coding regions) for expression quantification. Libraries with mapping rate < 40% were excluded from the subsequent analyses. With a consistent set of homologs, expression was quantified in transcripts per million (TPM) and thus was comparable across species. Only homologs expressed (TPM > 1) in > 10% of samples were retained for subsequent association analysis. For each taxon, we calculated the average expression of homologs among the replicates.

## Cross-species genomic and transcriptomic association analyses

To tackle the genetic variation of the rhizosphere N traits amongst the assayed species, we modeled the trait variation with the functionality and regulation of each homolog separately. First, we tested the impact of loss-of-function (LoF) of each homolog on trait variation with the following model:

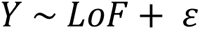

Where Y stands for average trait values, LoF denotes the binary scores that predict potential functional loss of the homolog, and 𝜀 is the residual.

Similarly, an association model was applied to test for the impact expression/dosage changes on trait evolution:

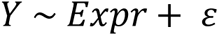

Where Y is the trait vector, Expr stands for the expression of each homolog in log_2_-scale and 𝜀 is the residual.

A matrix of shared branch length among species in the species tree was included in the models as a random effect to control for species relatedness (shared macroevolutionary history) as proposed previously (Lynch, 1991). However, potentially due to the star-like phylogeny of the assayed species or the phylogenetically independent evolution of rhizosphere N dynamics, including this random effect term had little impact on the test results (Supplementary Figure S2). Hence, we focus on the results of the fixed effect model for simplicity.

## Candidate genes

The moderate sample size (N = 36) compromised the power of the above trait-by-homolog association models at a genome wide scale. Hence, we focused on an *a priori* set of 188 candidate genes previously shown to affect BNI compound synthesis and transport as well as N uptake and mobilization in maize and sorghum (Supplementary Table S4) (Léran *et al*., 2014; Tesfamariam *et al*., 2014; Widhalm & Rhodes, 2016; Wang *et al*., 2020; Pan *et al*., 2021; Otaka *et al*., 2022; Petroli *et al*., 2023).

## Results

### The evolution of distinct rhizosphere N cycle management strategies

We measured the potential nitrification rate (PNR), NO_3_^-^ production and loss in the rhizosphere system of various Andropogoneae species grown on US corn belt soil during their active vegetative growth stage (Figure 2a). Significant variations were observed across species in all three rhizosphere N traits (Table 1). However, this variation did not show a clear association with the neutral phylogeny, indicating independent trait evolution (Figure 2b). A rhizosphere system with higher nitrification potential accumulates more NO_3_^-^ (Figure 2d) while this does not necessarily translate to higher NO_3_^-^ leaching (Figure 2c). This suggests that some species may be able to retain or utilize NO_3_^-^ differently. Comparing the potential nitrification rate against the NO_3_^-^ loss (Figure 2c), we showcase that three distinct N cycle management strategies within the rhizosphere systems have evolved in the Andropogoneae tribe. Interestingly, the rhizosphere soil of all examined annual species, irrespective of domestication status (*Sorghum bicolor*, *Zea mays*, and *Thelepogon elegans*), exhibited pronounced nitrification inhibition (potential nitrification was reduced by 25% or more compared to bulk soil potential). These species produced and lost the least NO_3_^-^, thereby conserving N within the system (Figure 2c). Vetiver, *Chrysopogon zizanioides*, is the only perennial species that demonstrated similar N conservation traits. We categorize these species as “**Conservationists**.” In contrast, most perennial grasses exhibited no significant control over the nitrification rate in their rhizosphere, with some even *enhancing* nitrification markedly (nitrification was elevated by 25% or more compared to bulk soil potential). Seven of these nitrification-enhancing species (*Elionorus tripsacoides*, *Themeda triandra*, *Urelytrum digitatum*, *Sorghastrum nutans*, *Schizachyrium scoparium*, *Anatherum virginicum* and *Schizachyrium microstachyum*) displayed increased NO_3_^-^ production and leaching (>50% of bulk soil NO_3_^-^ loss), as anticipated (Figure 2c). These species are referred to as “**Leachers**.” Intriguingly, another group of species (*Ischaemum rugosum*, *Miscanthus junceus*, *Pogonatherum paniceum* and *Andropogon gerardi*) enhanced rhizosphere nitrification while minimizing NO_3_^-^ loss from the system (<50% of bulk soil NO_3_^-^ loss) (Figure 2c). We term these species “**Nitrate Keepers**.”

**Figure 2.**
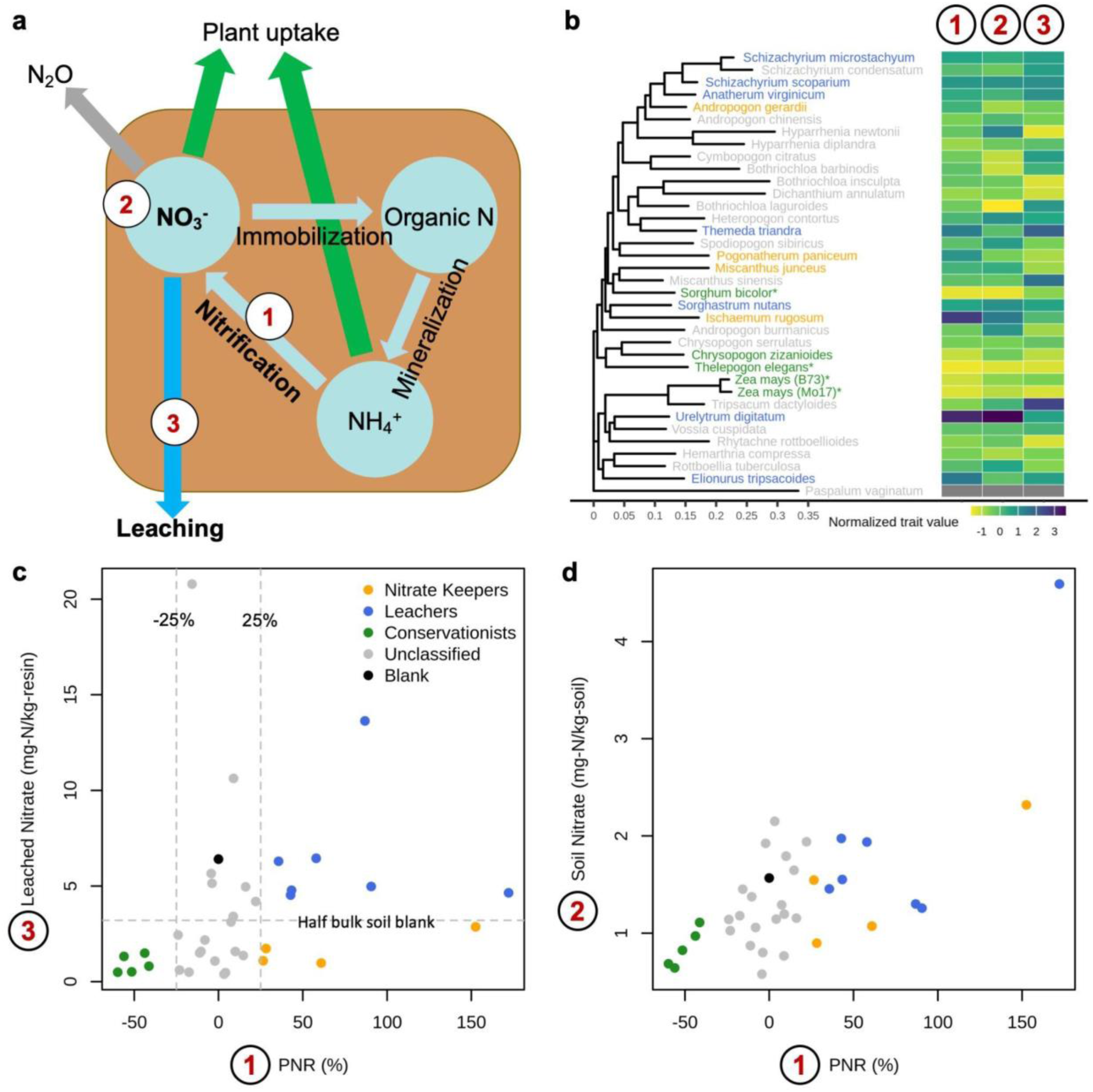
Soil N measurements and the classification of distinct rhizosphere N managing strategies. a. Illustration of the soil N cycle. Soil N exists as organic N, NH_4_ ^+^ and NO_3_^-^. Via mineralization processes, organic N is mobilized into NH_4_ ^+^. NH_4_ ^+^ can be nitrified into NO_3_^-^. Through immobilization processes, inorganic NO_3_^-^ turns back into organic N. Due to high mobility, NO_3_^-^ is more likely to escape the system, either through leaching or subjected to denitrification and turned to N_2_ and N_2_O. Given our focus on how diverse grass species manage their rhizosphere N budget, we measured (1) the potential nitrification rates (PNR) in different studied taxa relative to a bulk soil blank sample (in percentage), (2) soil nitrate concentrations (PPM) and (3) the amount of nitrate leached (PPM). **b.** Phylogenetically independent evolution of the rhizosphere N traits. Extremes of the measured rhizosphere N traits evolved multiple times across the Andropogoneae phylogeny. Trait values are shown after normalization ((x- mean(x))/sd(x)) as heatmap. Colors in tip labels refer to panel c. **c-d**. Scatter plots amongst the three measured N traits. The classification of distinct rhizosphere N managing strategies for different taxa is based on **c**. Relative to a bulk soil blank sample, “Conservationists” inhibit PNR by 25% or more and leach NO_3_^-^ by 50% or less; “Leachers” elevated PNR by 25% or more and leach NO_3_^-^ by 50% or more, and “Nitrate keepers” increase PNR by 50% or more but lost NO_3_^-^ by 50% or less.

**Table 1.**
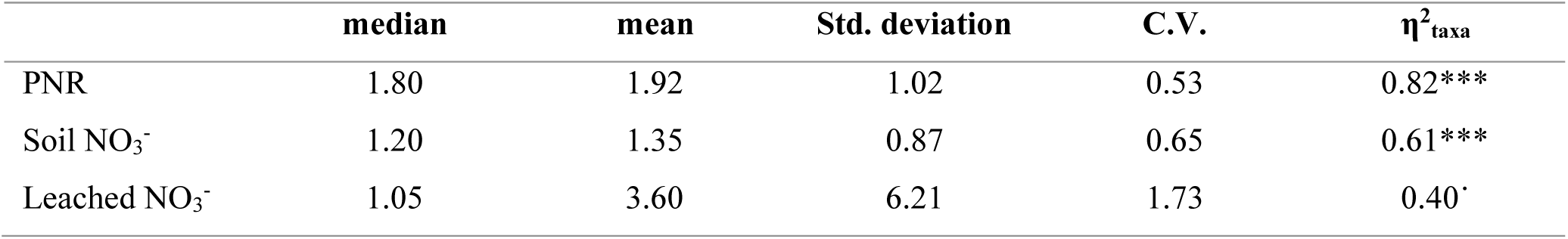
Descriptive statistics for soil N measurements. Potential nitrification rate (PNR) in mg N kg^-1^ hr^-1^, soil NO_3_^-^ concentration (mg N kg^-1^; PPM) and leached NO_3_^-^ concentration (mg N kg^-1^; PPM). The sample size equals 112 which is composed of 37 taxa each with 1-3 replicates. The variance explained by taxa (η^2^) are calculated based on the analysis of variance (ANOVA) using log-transformed trait data as following: η^2^ = SS /SS . The significance of the taxa term in ANOVA is denoted as following: *** p < 0.001; p < 0.1.

## Ecological selection on rhizosphere N cycle

To understand the potential ecological selection forces that drive the evolution of distinct rhizosphere N management strategies in different species, we associated the biogeography and habitat environmental conditions of diverse Andropogoneae species with their rhizosphere N traits. The range records of different species were retrieved from the Botanical Information and Ecology Network (BIEN). Using all the documented occurrence coordinates, we collected a total of 291 environmental features (see materials and methods) and summarized the environmental variation with 20 principal components (envPCs) which account for >80% of the total environmental variance (Supplementary Figure S3). Association analysis revealed five envPCs that significantly explain the rhizosphere N trait variation (Figure 3a). Several soil characteristics (e.g. acidity, nutrient composition, temperature, particle size…etc) and climatic factors (e.g. isothermality) loaded heavily on the significant rhizosphere N-associated envPCs (Supplementary Figure S4). “Leachers” originate from areas with drier and sandier soil with high leaching potential (Figure 3b-d). “Conservationists” with nitrification inhibitory effects are adapted to less acidic (Figure 3e and j), more fertile (Figure 3f and g) soil where nitrifying microbes may be more active. We also observed slightly higher soil isothermality (i.e. diurnal changes are closer to seasonal difference) in the habitat range of the “Conservationists” (Figure 3i). The “Nitrate keepers” originally live on potassium (K)-rich soil (Figure 3h). This result implied that the diverse rhizosphere N management strategies in the Andropogoneae tribe may evolve as potential adaptive responses in distinct ecological niches.

**Figure 3.**
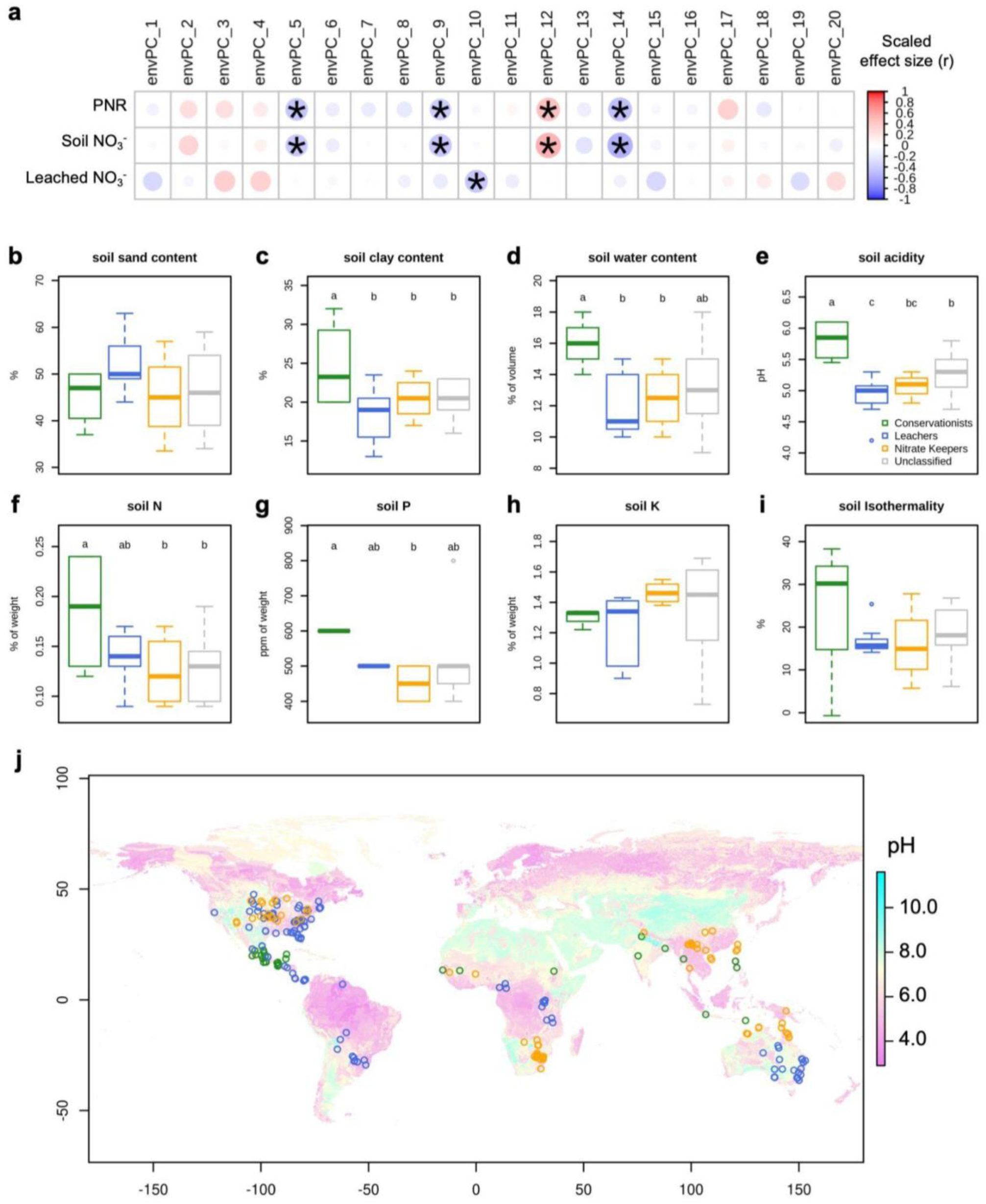
The association of rhizosphere N traits and the native habitat environments. a. A summary of the association analyses between the habitat environments of different species and their rhizosphere N traits. The habitat environments of different species are summarized with a principal component analysis (PCA) of 282 environmental features, where the top 20 envPCs explained >80% of total variance. A list of the environmental features and the loading of the PCA is available in Supplementary table S2. We test the association of each envPC with each rhizosphere N trait. Each cell indicates an independent test. The color and the circle size denote the effect of the association (Pearson’s correlation coefficient). Asterisks denote statistical significance (p < 0.05). Five envPCs significantly co-vary with the rhizosphere N trait variation. We select a subset of the top contributing environmental features to these significant envPCs and visualize the difference between species with different rhizosphere N managing strategies. These include **b.** soil sand content, **c.** soil clay content, **d.** soil water content, **e.** soil pH, **f.** soil N, **g.** soil phosphate (P), **h.** soil potassium (K) and **i.** soil isothermality. Small case letters denote statistical grouping based on Fisher’s Least Significant Difference test. **j**. A map of global soil acidity (pH) with species occurrence records (empty dots) overlaid. Occurrence data are retrieved for each studied Andropgoneae species from the BIEN database. Species with >20 occurrence records are down-sampled to 20 records for visual clarity. Dot colors follow the species group colors denoted in panel e. Both leacher and nitrate keeper species (blue and orange) showed occurrence in more acidic environments.

## Genetic basis of the variation in rhizosphere N cycle

Next, we explored how the potential loss of function and regulatory evolution of the homologs affect the rhizosphere N cycle. With a linear modeling framework, we conducted a total of nine sets of cross-species association analyses between the three rhizosphere N traits (PNR, soil NO_3_^-^ and NO_3_^-^ loss) and three genetic factors (loss of function prediction, leaf expression and root expression). The moderate number of species assayed in this study limits the statistical power of the association models. Therefore, we focus our gene discovery on a set of 188 candidates (Supplementary Table S4) identified from previous works (Léran *et al*., 2014; Tesfamariam *et al*., 2014; Widhalm & Rhodes, 2016; Wang *et al*., 2020; Pan *et al*., 2021; Otaka *et al*., 2022; Petroli *et al*., 2023). We aim to test for shared evolution signatures in multiple species that evolved similar rhizosphere N managing strategies. Compared to a random set of homologs, *a priori* candidate homologs are enriched for significant association at the p-value threshold of 0.01, particularly between root expression profiles and rhizosphere N traits (Supplementary Figure S5). We identified 13 homologs that are key to the variation of rhizosphere N cycle in diverse species (Figure 4a and Supplementary Figure S6; Supplementary Table S5).

**Figure 4.**
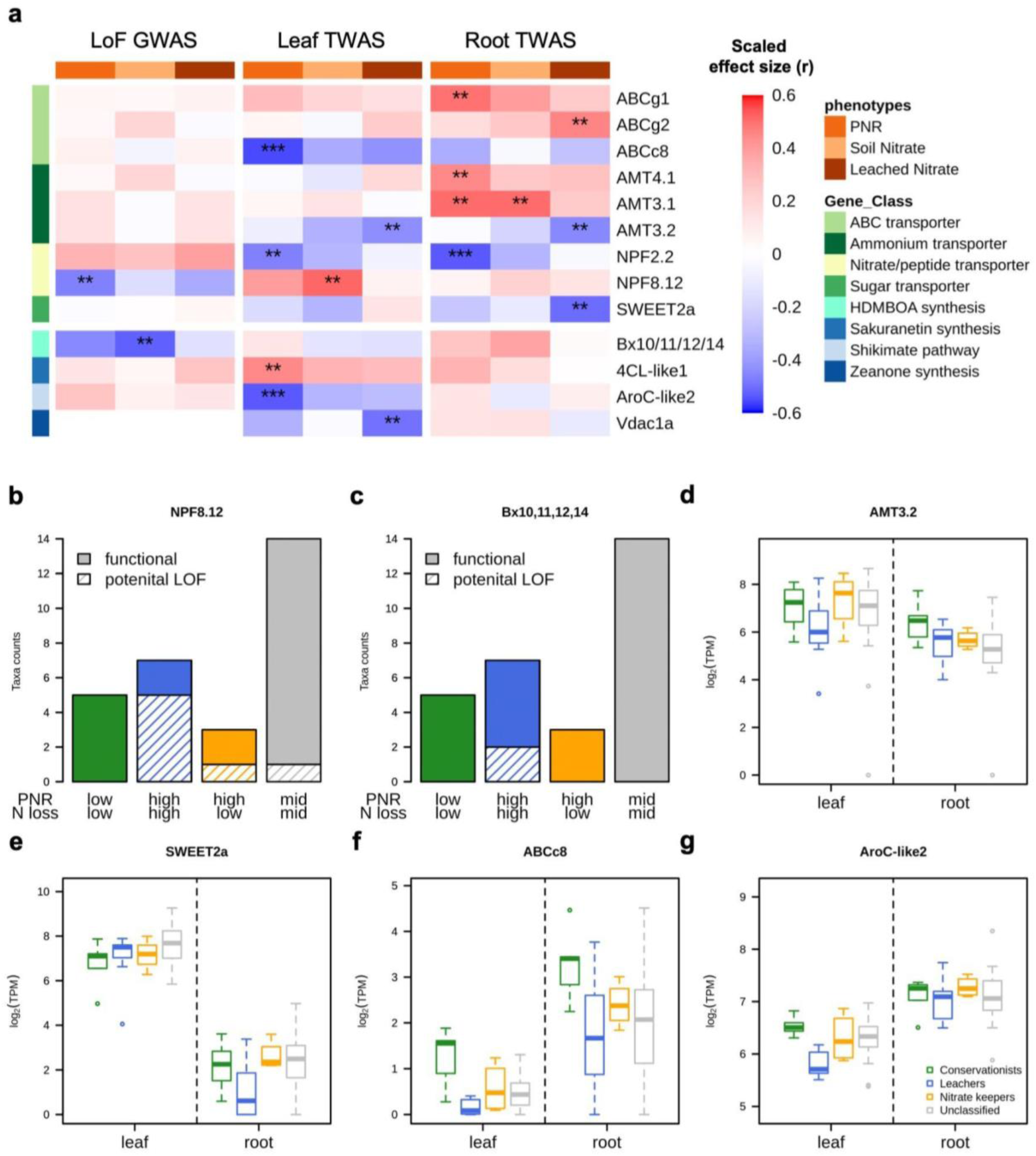
Genes underlying the rhizosphere N trait variation. a. A summary of the significant association between different genetic factors (i.e. LoF prediction and expression estimates) and the rhizosphere N traits across species for the candidate homologs. For each homolog, there are nine association tests (three genetic factors by three rhizosphere N traits) as denoted by the columns. Each row is a homolog that exhibits significant association in at least one trait-genetic combination. Row annotation colors indicate the gene classes each significant homolog belongs to. The cell colors denote the strength and direction of the association (Pearson’s correlation coefficient, r). Stars denote statistical significance (**: p < 0.01; ***: p < 0.001). The same test summary for all tested homologs is available as Supplementary Figure S6. **b. & c.** Bar plots demonstrating the occurrence of potential LoF in different species groups with distinct rhizosphere N traits for the homologs of NPF8.12 and Bx10,11,12,14 respectively. Bar heights indicate the count of taxa. Colors denote the species groups. Solid bars represent the functional copies while slashed bars indicate potential loss of function. **d-g.** Homolog expression profiles in different tissues (left: leaf and right: root) in different species groups with distinct rhizosphere N traits (colors) for a subset of candidates: **d.** AMT3.2, **e.** SWEET2a, **f.** ABCc8, and **g.** AroC-like2. The remaining significant homolog- trait associations are visualized in Supplementary Figure S7.

The significant candidates mostly encode transporter proteins, which include three ammonium transporters (AMTs), two nitrate/peptide transporter family proteins (NPFs), one sugar transporters (SWEETs) and three ABC-type transporters (Figure 4a). The potential loss of function of ZmNPF8.12 homolog is significantly associated with higher nitrification rate. >70% of the Leachers, which produced and lost the most NO_3_^-^, do not have a functional copy of this homolog (Figure 4b). In line with this, we detected a signature of relatively constrained evolution for ZmNPF8.12 homologs in the Conservationists species (codeml-branch model; likelihood ratio = 6.58, p = 0.01). Leaf expression of the ZmNPF2.2/2.3 homolog appears to be important for N conservation (Supplementary Figure S7). The homolog of ZmAMT3.2 exhibits relatively high expression in the roots of the Conservationists species which exhibit strong nitrification inhibitory effects (Figure 4d). Root expression of the ZmSWEET2a homolog is associated with lower NO_3_^-^ leaching, with low expression in the roots of the leacher species (Figure 4e). In addition to the nutrient transporters, the expression of the ZmABCc8 homolog is negatively correlated with rhizosphere nitrification rate (Figure 4f).

Four homologs of the genes involved in BNI metabolite biosynthesis were detected, one of which is found in the benzoxazinoid biosynthetic pathway (Figure 4a). HDMBOA, an important BNI metabolite in maize (Otaka *et al*., 2022), is a benzoxazinoid derivative (2-hydroxy-4,7-dimethoxy- 1,4-bezoxazin-3-one). Potential loss of function of the ZmBx10/11/12/14 homolog, which catalyzes the synthesis of the storage form HDMBOA-glucoside (Otaka *et al*., 2022), is associated with increased production of rhizosphere NO_3_^-^ (Figure 4a): we only observe putative loss of function events in two Leacher species (Figure 4c). In addition, we also observe a significant negative correlation between the NO_3_^-^ loss rate and the leaf expression of ZmVdac1a homolog. ZmVdac1a encodes an isochorismate synthase which is potentially important for the synthesis of zeanone, another BNI metabolite in maize (Otaka *et al*., 2022). The leaf expression of another homolog encoding a ZmAroC-like protein correlates negatively with soil nitrification: higher expression is observed in the conservationist species (Figure 4g).

Despite the limited statistical power, top outliers with the strongest association signatures among all tested homologs may also be reliable candidates, in addition to the *a priori* ones. We identified the top five candidates in each of the nine association tests and collected a union set of 40 homologs (Supplementary Table S5). These homologs are enriched for biological functions related to defense responses (GO:0050832, GO:0042742 and GO:0031640) as well as cell wall and polysaccharide catabolic processes (GO:0006032, GO:0016998 and GO:0000272) (Supplementary Table S6). The potential loss of function in a homolog encoding a knottin scorpion toxin-like protein and low root expression of the ZmSod2 (superoxide dismutase) homolog are linked to elevated NO_3_^-^ loss. Additionally, increased expression of several homologs with putative chitinase, cellulase, and laccase activities correlates with higher NO_3_^-^ loss. These findings highlight the significance of plant-microbe interactions in the rhizosphere for soil N dynamics.

## Discussion

In this study, we present the first systematic screening for rhizosphere N dynamics in dozens of grass species with wide adaptive ranges, identifying three distinct patterns of N-cycling (Figure 5). BNI capacity, in combination with a “more ammonium” management, was proposed as one important trait for soil N sustainability (Subbarao & Searchinger, 2021). The documentation and success of BNI capacities in perennial grasses in tropical savannas (Subbarao *et al*., 2007a; Srikanthasamy *et al*., 2018) and pasture systems (Baruch *et al*., 1985; Rossiter-Rachor *et al*., 2009) gave rise to the hypothesis that some Andropogoneae species may present enhanced BNI capacity and potentially favorable rhizosphere N traits that are not observed in the standing genetic variation of domesticated annual maize. However, we observe little evidence supporting this hypothesis. Despite substantial variation across diverse Andropogoneae species, only one wild annual grass, *Thelepogon elegans*, exhibits slightly higher BNI capacity and N conservation than the domesticated annuals in our experiment. Both annual crops assayed in this experiment, maize and sorghum, are already good N Conservationists. This result undermines the leverage of tribe-level phylogenetic diversity for BNI improvement in maize. However, the systematic assays of diverse grasses allowed unprecedented biogeographic and cross-species genomic/transcriptomic analyses on the emergence and evolution of rhizosphere N cycles (Figure 5).

**Figure 5.**
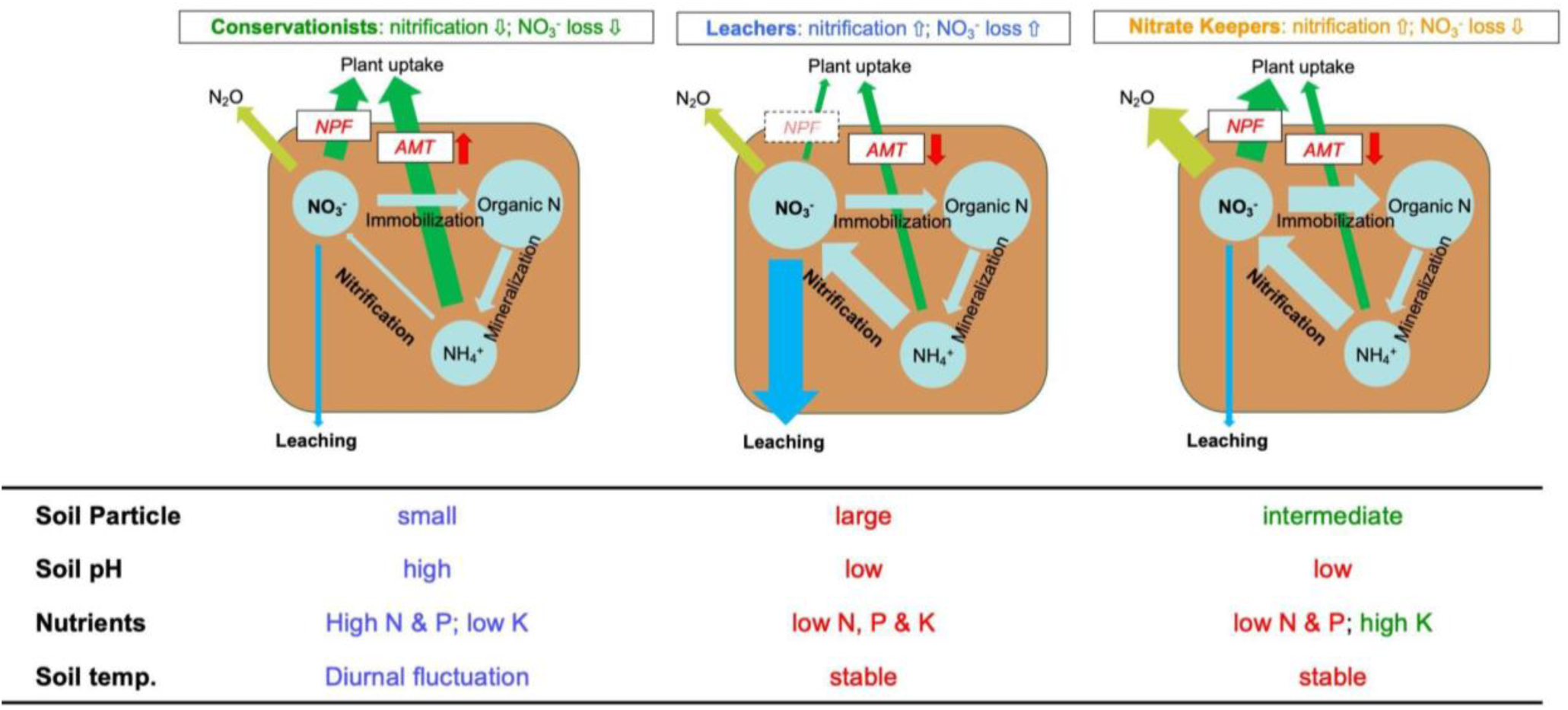
**Schematic illustration of diverse rhizosphere N cycles in Andropogoneae.** This figure illustrates the hypothetical rhizosphere N cycles of three species groups with distinct rhizosphere N managing strategies evolved in the Andropogoneae tribe: Conservationists, Leachers, and Nitrate Keepers. The Conservationists inhibit nitrification and potentially uptake the conserved NH_4_^+^ with the higher expression of ZmAMT3.2 homolog. The Leachers stimulate nitrification but are not able to utilize the converted NO_3_^-^ potentially due to the lack of functional ZmNPF8.12 nitrate transporter. In contrast, the Nitrate Keepers possess functional copies of ZmNPF8.12 and thus they lose less NO_3_^-^. Conservationists originate from more fertile, less acidic and moist soils. In contrast, the Leachers are from habitats with drainage and/or nutrient stresses. One key difference between the habitats of the Nitrate Keepers and the one of the Leachers is the drainage stress.

Three out of the four conservationist species exhibit annual life history, implicating potential benefit of such N conservation traits during the perennial-annual transition. We observed that the conservationist species originate from more fertile areas (high N and P; Figure 3f-g and Figure 5). In contrast to the hypothesis that low N availability drives the evolution of BNI (Lata *et al*., 2022), our result suggests the competition with nitrifiers for more available N may be the driving force for the emergence of BNI. Notably, in addition to N fertility, we also observed a significant association between BNI capacity and soil pH. Species with high BNI capacity grow on less acidic soil (Figure 3e and Figure 5). Independent evidence for the emergence of BNI capacity during the adaptation to alkali soils was also documented recently (Wang *et al*., 2023; Przybylska *et al*., 2024). We expect limited nitrifier populations in more acidic environments as the nitrification reaction is largely inhibited when soil pH is below 5 (Dancer *et al*., 1973). Hence, we hypothesize that BNI capacity and NH_4_^+^ conservation mechanisms in a few Andropogoneae grasses could emerge in competition with nitrifying bacteria during evolution. Nevertheless, maize field soil was used in our common garden experiment so it is also possible that the lack of nitrification inhibitory or even stimulating effects in some of the wild species may be mal-adaptive plasticity in response to the foreign soil and microbes. Multiple common garden experiments including distinct soil types or reciprocal transplantation between some species pairs will provide further insights into this hypothesis.

The discovery of Nitrate Keepers, which enhanced nitrification while conserving NO_3_^-^, is interesting. There is massive variation in the preference between NH_4_ ^+^ and NO_3_^-^ among plant species (Houlton *et al*., 2007; Kahmen *et al*., 2008; Boudsocq *et al*., 2012; Britto & Kronzucker, 2013), and such preference determines the fitness benefits and cost of BNI capacity (Boudsocq *et al*., 2012; Konaré *et al*., 2019). Stimulation of nitrification can be particularly beneficial for NO_3_^-^-preferring species in competition to NH_4_ ^+^-preferring plants. The example of two ecotypes with opposing effects on nitrification for *Hyparrhenia diplandra* (Lata *et al*., 2000) highlights the presence of environment-specific plant-plant and plant-microbe competition for different N-sources and the evolution of locally adapted soil N management strategies. Therefore, we speculate the “Nitrate Keepers” may exhibit a preferential uptake of NO_3_^-^ over NH ^+^ while further experiments are still needed.

In this study, each species was represented by only one genotype, which is a potential caveat: different ecotypes within a species might influence soil chemistry differently (e.g. (Lata *et al*., 2000). Examining within-species variation, while potentially important and insightful, was beyond the scope of the study, which was designed for broad interspecific differences exploration. We encourage future studies to follow up on the variation within a couple of the species or to include multiple genotypes per species in similar analyses.

This substantial cross-species variation in rhizosphere N dynamics underscores the importance of understanding the genetic underpinnings of these traits. The enrichment of transporter-encoding homologs, particularly N transporters, in our cross-species association analyses consistently highlights the relevance of soil N dynamics and plant preference. On the one hand, nitrification-inhibiting Conservationists have higher root expression of ammonium transporter (ZmAMT3.2), implicating their potential preference for conserved NH_4_^+^ (Figure 4d and Figure 5). On the other hand, most Nitrate Keepers exhibit lower root expression of ZmAMT3.2 (Figure 4d) but preserve a functional copy of the ZmNPF8.12 nitrate transporter homolog unlike the Leachers, implying their potential preference for NO_3_^-^ (Figure 4b and Figure 5) Interestingly, another ammonium transporter paralog, ZmAMT5, was identified in a recent GWAS study on maize BNI (Petroli *et al*., 2023). The independent discovery of association between BNI activity and AMT gene family supports the interaction between N uptake/preference and soil nitrification modification.

Besides N transporters, several homologs encoding ABC-type transporters were identified by our association analyses. A homolog of ABC type c transporter (ZmABCc8) showed higher expression in the nitrification-inhibiting Conservationist species. Maize ZmABCc8 and its ortholog in Arabidopsis were reported to be involved in the transport of anthocyanin (Goodman *et al*., 2004; Dean *et al*., 2022). Anthocyanin and other flavonoids play a role in copper chelation (Sarma *et al*., 1997), which is thought to be one nitrification inhibitory mechanism (Corrochano-Monsalve *et al*., 2021). Therefore, we hypothesized that the identified ABC type c transporter may be responsible for the exudation of BNI compounds and thus their expression correlates with the BNI capacity.

In addition to the transporters, we also detected significant association in a handful of homologs involved in the biosynthesis of known BNI compounds in maize and sorghum. The identification of homologs involved in the biosynthesis and the modification of chorismate highlight its central role in the synthesis of BNI metabolites. However, the wide variety of BNI compounds found in different BNI-species (Subbarao *et al*., 2007b, 2013; Pariasca Tanaka *et al*., 2010; Sun *et al*., 2016; Otaka *et al*., 2022) suggests the presence of multiple biochemical routes for the gain of BNI capacity. This poses a challenge for the identification of more genes with convergent functional/regulatory evolution.

The choice of US corn belt soil for the experiment makes our results directly relevant to modern maize production. Our systematic exploration of phylogenetic diversity in rhizosphere N dynamics offers insights into how and/or to which extent breeding efforts could reduce the environmental impact of maize agriculture. Fine-tuning the regulation of our candidate transporters may help optimize maize rhizosphere N balance. For instance, early expression of AMTs might make maize better fit to the “more-ammonium” management (Subbarao & Searchinger, 2021). The discovery of NO_3_^-^ keeping strategies also highlight the potential of early and efficient nitrate uptake. Since maize evolved as one of the conservationists in the tribe, favorable alleles might have emerged and segregated as natural variation within maize. The field will need more studies like the recent GWAS (Petroli *et al*., 2023) to tap the genetic diversity within maize and more exploration in other wild annual *Zea* species not included in this study. Beyond genetic improvement for N uptake and conservation during the growing season of the annual crops, the integration of cover crops and intercropping into the agricultural system can also be crucial, if not more so, for mitigating early spring nitrogen loss (Parkin *et al*., 2016; Zhang *et al*., 2023; Rogovska *et al*., 2023).

## Acknowledgement

This project is supported by the NSF PanAnd Grant Award #1822330 and USDA-ARS. Thank you Corteva and INARI for genomic data generation. Thank you Sara Miller, Zong-Yan Liu, Sam Herr and Elad Oren for support on sample collection during the experiment. Graph design is facilitated by resources on Flaticon.com, D-ALLE, and BioRender.com.

## Author contribution

S-KH, BDE, SSR, ADK, CR, and ESB designed the research. TMA-E and EAK collected the materials of the research. S-KH, NL, TL, BDE and AJH performed the experiment and collected the data. S-KH, BDE, GC-N, AJS and COH analyzed the data. S-KH, BDE, GC-N, SSR, JOO-R, EAK and ESB were involved in result interpretations. S-KH drafted the manuscript and all other authors edited the manuscript.

## Data availability

All data, source scripts, analytical notebooks and intermediate outputs are available at (https://bitbucket.org/bucklerlab/p_evolBNI_publication/src/main/). The raw sequencing read data in this study have been submitted to the NCBI BioProject database (https://www.ncbi.nlm.nih.gov/bioproject/) under accession number PRJEB50280 (genomic long read), PRJNA543119 (genomic short reads) and PRJNA1119410 (RNA short reads). Short read assemblies will be made available on Zenodo upon publication (10.5281/zenodo.11222298).

## Reference

1. Abalos D, van Groenigen JW, De Deyn GB. 2018. What plant functional traits can reduce nitrous oxide emissions from intensively managed grasslands? Global change biology 24: e248– e258.

2. Bachle S, Zaricor M, Griffith D, Qui F, Still CJ, Ungerer MC, Nippert JB. 2022. Physiological responses to drought stress and recovery reflect differences in leaf function and anatomy among grass lineages. bioRxiv: 2022.07.30.502130.

3. Baruch Z, Ludlow MM, Davis R. 1985. Photosynthetic responses of native and introduced C4 grasses from Venezuelan savannas. Oecologia 67: 388–393.

4. Bender RR, Haegele JW, Ruffo ML, Below FE. 2013. Nutrient uptake, partitioning, and remobilization in modern, transgenic insect-protected maize hybrids. Agronomy journal 105: 161–170.

5. Billen G, Garnier J, Lassaletta L. 2013. The nitrogen cascade from agricultural soils to the sea: modelling nitrogen transfers at regional watershed and global scales. Philosophical transactions of the Royal Society of London. Series B, Biological sciences 368: 20130123.

6. Boudsocq S, Niboyet A, Lata JC, Raynaud X, Loeuille N, Mathieu J, Blouin M, Abbadie L, Barot S. 2012. Plant preference for ammonium versus nitrate: a neglected determinant of ecosystem functioning? The American naturalist 180: 60–69.

7. Bremner JM, Blackmer AM. 1978. Nitrous oxide: emission from soils during nitrification of fertilizer nitrogen. Science 199: 295–296.

8. Britto DT, Kronzucker HJ. 2013. Ecological significance and complexity of N-source preference in plants. Annals of botany 112: 957–963.

9. Byrnes RC, Nùñez J, Arenas L, Rao I, Trujillo C, Alvarez C, Arango J, Rasche F, Chirinda N. 2017. Biological nitrification inhibition by Brachiaria grasses mitigates soil nitrous oxide emissions from bovine urine patches. Soil biology & biochemistry 107: 156–163.

10. Corrochano-Monsalve M, González-Murua C, Bozal-Leorri A, Lezama L, Artetxe B. 2021. Mechanism of action of nitrification inhibitors based on dimethylpyrazole: A matter of chelation. The Science of the total environment 752: 141885.

11. Cowan MF, Blomstedt CK, Norton SL, Henry RJ, Møller BL, Gleadow R. 2020. Crop wild relatives as a genetic resource for generating low-cyanide, drought-tolerant Sorghum. Environmental and experimental botany 169: 103884.

12. Dancer WS, Peterson LA, Chesters G. 1973. Ammonification and nitrification of N as influenced by soil pH and previous N treatments. Soil Science Society of America journal. Soil Science Society of America 37: 67–69.

13. Dean JV, Willis M, Shaban L. 2022. Transport of acylated anthocyanins by the Arabidopsis ATP-binding cassette transporters AtABCC1, AtABCC2, and AtABCC14. Physiologia plantarum 174: e13780.

14. Doane TA, Horwáth WR. 2003. Spectrophotometric Determination of Nitrate with a Single Reagent. Analytical letters 36: 2713–2722.

15. EPA. 2023. Inventory of U.S. Greenhouse Gas Emissions and Sinks: 1990-2021.

16. Esteban R, Ariz I, Cruz C, Moran JF. 2016. Review: Mechanisms of ammonium toxicity and the quest for tolerance. Plant science: an international journal of experimental plant biology 248: 92–101.

17. Estep MC, McKain MR, Vela Diaz D, Zhong J, Hodge JG, Hodkinson TR, Layton DJ, Malcomber ST, Pasquet R, Kellogg EA. 2014. Allopolyploidy, diversification, and the Miocene grassland expansion. Proceedings of the National Academy of Sciences of the United States of America 111: 15149–15154.

18. Fick SE, Hijmans RJ. 2017. WorldClim 2: new 1-km spatial resolution climate surfaces for global land areas. International Journal of Climatology 37: 4302–4315.

19. GAEZ v4 Data Portal. Goodman CD, Casati P, Walbot V. 2004. A multidrug resistance-associated protein involved in anthocyanin transport in Zea mays. The Plant cell 16: 1812–1826.

20. Hartman MD, Burnham M, Parton WJ, Finzi A, DeLucia EH, Yang WH. 2022. In silico evaluation of plant nitrification suppression effects on agroecosystem nitrogen loss. Ecosphere 13.

21. Hart SC, Stark JM, Davidson EA, Firestone MK. 1994. Nitrogen Mineralization, Immobilization, and Nitrification. In: SSSA Book Series. Methods of Soil Analysis. Madison, WI, USA: Soil Science Society of America, 985–1018.

22. Hijmans RJ, Bivand R, Pebesma E, Sumner MD. 2023. terra: R package for spatial data handling. Github.

23. Houlton BZ, Sigman DM, Schuur EAG, Hedin LO. 2007. A climate-driven switch in plant nitrogen acquisition within tropical forest communities. Proceedings of the National Academy of Sciences of the United States of America 104: 8902–8906.

24. Hufford MB, Seetharam AS, Woodhouse MR, Chougule KM, Ou S, Liu J, Ricci WA, Guo T, Olson A, Qiu Y, et al. 2021. De novo assembly, annotation, and comparative analysis of 26 diverse maize genomes. Science 373: 655–662.

25. Husson F, Josse J. 2023. *missMDA — Handling Missing Values with Multivariate Data Analysis*. Github.

26. Husson F, Josse J, Le S, Mazet J. 2023. *FactoMineR: Package FactoMineR*. Github.

27. Kahmen A, Wanek W, Buchmann N. 2008. Foliar delta(15)N values characterize soil N cycling and reflect nitrate or ammonium preference of plants along a temperate grassland gradient. Oecologia 156: 861–870.

28. Katoh K, Misawa K, Kuma K-I, Miyata T. 2002. MAFFT: a novel method for rapid multiple sequence alignment based on fast Fourier transform. Nucleic acids research 30: 3059–3066.

29. Khush GS. 2001. Green revolution: the way forward. Nature reviews. Genetics 2: 815–822.

30. Konaré S, Boudsocq S, Gignoux J, Lata J-C, Raynaud X, Barot S. 2019. Effects of Mineral Nitrogen Partitioning on Tree–Grass Coexistence in West African Savannas. Ecosystems 22: 1676–1690.

31. Kosola KR, Eller MS, Dohleman FG, Olmedo-Pico L, Bernhard B, Winans E, Barten TJ, Brzostowski L, Murphy LR, Gu C, et al. 2023. Short-stature and tall maize hybrids have a similar yield response to split-rate vs. pre-plant N applications, but differ in biomass and nitrogen partitioning. Field crops research 295: 108880.

32. Lata JC, Guillaume K, Degrange V, Abbadie L, Lensi R. 2000. Relationships between root density of the African grass Hyparrhenia diplandra and nitrification at the decimetric scale: an inhibition-stimulation balance hypothesis. Proceedings. Biological sciences / The Royal Society 267: 595–600.

33. Lata J-C, Le Roux X, Koffi KF, Yé L, Srikanthasamy T, Konaré S, Barot S. 2022. The causes of the selection of biological nitrification inhibition (BNI) in relation to ecosystem functioning and a research agenda to explore them. Biology and fertility of soils 58: 207–224.

34. Lembrechts JJ, van den Hoogen J, Aalto J, Ashcroft MB, De Frenne P, Kemppinen J, Kopecký M, Luoto M, Maclean IMD, Crowther TW, et al. 2022. Global maps of soil temperature. Global change biology 28: 3110–3144.

35. Léran S, Varala K, Boyer J-C, Chiurazzi M, Crawford N, Daniel-Vedele F, David L, Dickstein R, Fernandez E, Forde B, et al. 2014. A unified nomenclature of NITRATE TRANSPORTER 1/PEPTIDE TRANSPORTER family members in plants. Trends in plant science 19: 5–9.

36. Li H. 2023. Protein-to-genome alignment with miniprot. Bioinformatics 39.

37. Li D, Liu C-M, Luo R, Sadakane K, Lam T-W. 2015.MEGAHIT: an ultra-fast single-node solution for large and complex metagenomics assembly via succinct de Bruijn graph. Bioinformatics 31: 1674–1676.

38. Lu C, Yu Z, Zhang J, Cao P, Tian H, Nevison C. 2022. Century-long changes and drivers of soil nitrous oxide (N2 O) emissions across the contiguous United States. Global change biology 28: 2505–2524.

39. Lynch M. 1991. METHODS FOR THE ANALYSIS OF COMPARATIVE DATA IN EVOLUTIONARY BIOLOGY. Evolution; international journal of organic evolution 45: 1065–1080.

40. Maitner B. 2023. *RBIEN: Tools for accessing the Botanical Information and Ecology Network (BIEN) database*. Github.

41. McCormick RF, Truong SK, Sreedasyam A, Jenkins J, Shu S, Sims D, Kennedy M, Amirebrahimi M, Weers BD, McKinley B, et al. 2018. The Sorghum bicolor reference genome: improved assembly, gene annotations, a transcriptome atlas, and signatures of genome organization. The Plant journal: for cell and molecular biology 93: 338–354.

42. McNear DH Jr. 2013. The Rhizosphere - Roots, Soil and Everything In Between. Nature Education Knowledge 4: 1.

43. Miranda KM, Espey MG, Wink DA. 2001. A rapid, simple spectrophotometric method for simultaneous detection of nitrate and nitrite. Nitric oxide: biology and chemistry / official journal of the Nitric Oxide Society 5: 62–71.

44. Mitros T, Session AM, James BT, Wu GA, Belaffif MB, Clark LV, Shu S, Dong H, Barling A, Holmes JR, et al. 2020. Genome biology of the paleotetraploid perennial biomass crop Miscanthus. Nature communications 11: 5442.

45. Moore NA, Camac JS, Morgan JW. 2019. Effects of drought and fire on resprouting capacity of 52 temperate Australian perennial native grasses. The New phytologist 221: 1424–1433.

46. Nuñez J, Arevalo A, Karwat H, Egenolf K, Miles J, Chirinda N, Cadisch G, Rasche F, Rao I, Subbarao G, et al. 2018. Biological nitrification inhibition activity in a soil-grown biparental population of the forage grass, Brachiaria humidicola. Plant and soil 426: 401–411.

47. Otaka J, Subbarao GV, Ono H, Yoshihashi T. 2022. Biological nitrification inhibition in maize—isolation and identification of hydrophobic inhibitors from root exudates. Biology and fertility of soils 58: 251–264.

48. Pan Z, Bajsa-Hirschel J, Vaughn JN, Rimando AM, Baerson SR, Duke SO. 2021. In vivo assembly of the sorgoleone biosynthetic pathway and its impact on agroinfiltrated leaves of Nicotiana benthamiana. The New phytologist 230: 683–697.

49. Pariasca Tanaka J, Nardi P, Wissuwa M. 2010. Nitrification inhibition activity, a novel trait in root exudates of rice. AoB plants 2010: lq014.

50. Parkin TB, Kaspar TC, Jaynes DB, Moorman TB. 2016. Rye cover crop effects on direct and indirect nitrous oxide emissions. Soil Science Society of America journal. Soil Science Society of America 80: 1551–1559.

51. Patro R, Duggal G, Love MI, Irizarry RA, Kingsford C. 2017. Salmon provides fast and bias- aware quantification of transcript expression. Nature methods 14: 417–419.

52. Petroli CD, Subbarao GV, Burgueño JA, Yoshihashi T, Li H, Franco Duran J, Pixley KV. 2023. Genetic variation among elite inbred lines suggests potential to breed for BNI-capacity in maize. Scientific reports 13: 13422.

53. Przybylska MS, Violle C, Vile D, Scheepens JF, Munoz F, Tenllado Á, Vinyeta M, Le Roux X, Vasseur F. 2024. Can plants build their niche through modulation of soil microbial activities linked with nitrogen cycling? A test with Arabidopsis thaliana. The New phytologist.

54. Rabalais NN, Turner RE. 2019. Gulf of Mexico hypoxia: Past, present, and future. Limnology and Oceanography Bulletin 28: 117–124.

55. Rogovska N, O’Brien PL, Malone R, Emmett B, Kovar JL, Jaynes D, Kaspar T, Moorman TB, Kyveryga P. 2023. Long-term conservation practices reduce nitrate leaching while maintaining yields in tile-drained Midwestern soils. Agricultural water management 288: 108481.

56. Rossiter-Rachor NA, Setterfield SA, Douglas MM, Hutley LB, Cook GD, Schmidt S. 2009. Invasive Andropogon gayanus (gamba grass) is an ecosystem transformer of nitrogen relations in Australian savanna. Ecological applications: a publication of the Ecological Society of America 19: 1546–1560.

57. Ross CW, Prihodko L, Anchang J, Kumar S, Ji W, Hanan NP. 2018. HYSOGs250m, global gridded hydrologic soil groups for curve-number-based runoff modeling. Scientific data 5: 180091.

58. Sarma AD, Sreelakshmi Y, Sharma R. 1997. Antioxidant ability of anthocyanins against ascorbic acid oxidation. Phytochemistry 45: 671–674.

59. Schlesinger WH. 2009. On the fate of anthropogenic nitrogen. Proceedings of the National Academy of Sciences of the United States of America 106: 203–208.

60. Schulz AJ, Zhai J, AuBuchon-Elder T, El-Walid M, Ferebee TH, Gilmore EH, Hufford MB, Johnson LC, Kellogg EA, La T, et al. 2023. Fishing for a reelGene: evaluating gene models with evolution and machine learning. bioRxiv: 2023.09.19.558246.

61. Shangguan W, Dai Y, Duan Q, Liu B, Yuan H. 2014. A global soil data set for earth system modeling. Journal of advances in modeling earth systems 6: 249–263.

62. Smith C, Hill AK, Torrente-Murciano L. 2020. Current and future role of Haber–Bosch ammonia in a carbon-free energy landscape. Energy & environmental science 13: 331–344.

63. Srikanthasamy T, Leloup J, N’Dri AB, Barot S, Gervaix J, Koné AW, Koffi KF, Le Roux X, Raynaud X, Lata J-C. 2018. Contrasting effects of grasses and trees on microbial N-cycling in an African humid savanna. Soil biology & biochemistry 117: 153–163.

64. Stamatakis A. 2014. RAxML version 8: a tool for phylogenetic analysis and post-analysis of large phylogenies. Bioinformatics 30: 1312–1313.

65. Subbarao GV, Kishii M, Bozal-Leorri A, Ortiz-Monasterio I, Gao X, Ibba MI, Karwat H, Gonzalez-Moro MB, Gonzalez-Murua C, Yoshihashi T, et al. 2021. Enlisting wild grass genes to combat nitrification in wheat farming: A nature-based solution. Proceedings of the National Academy of Sciences of the United States of America 118: e2106595118.

66. Subbarao GV, Nakahara K, Hurtado MP, Ono H, Moreta DE, Salcedo AF, Yoshihashi AT, Ishikawa T, Ishitani M, Ohnishi-Kameyama M, et al. 2009. Evidence for biological nitrification inhibition in Brachiaria pastures. Proceedings of the National Academy of Sciences of the United States of America 106: 17302–17307.

67. Subbarao GV, Nakahara K, Ishikawa T, Ono H, Yoshida M, Yoshihashi T, Zhu Y, Zakir HAKM, Deshpande SP, Hash CT, et al. 2013. Biological nitrification inhibition (BNI) activity in sorghum and its characterization. Plant and soil 366: 243–259.

68. Subbarao GV, Rondon M, Ito O, Ishikawa T, Rao IM, Nakahara K, Lascano C, Berry WL. 2007a. Biological nitrification inhibition (BNI)—is it a widespread phenomenon? Plant and soil 294: 5–18.

69. Subbarao GV, Searchinger TD. 2021. A ‘more ammonium solution’ to mitigate nitrogen pollution and boost crop yields. Proceedings of the National Academy of Sciences 118: e2107576118.

70. Subbarao GV, Tomohiro B, Masahiro K, Osamu I, Samejima H, Wang HY, Pearse SJ, Gopalakrishnan S, Nakahara K, Zakir Hossain AKM, et al. 2007b. Can biological nitrification inhibition (BNI) genes from perennial Leymus racemosus (Triticeae) combat nitrification in wheat farming? Plant and soil 299: 55–64.

71. Subbarao GV, Wang HY, Ito O, Nakahara K, Berry WL. 2007c. NH4+triggers the synthesis and release of biological nitrification inhibition compounds in Brachiaria humidicola roots. Plant and soil 290: 245–257.

72. Sun L, Lu Y, Yu F, Kronzucker HJ, Shi W. 2016. Biological nitrification inhibition by rice root exudates and its relationship with nitrogen-use efficiency. The New phytologist 212: 646– 656.

73. Sun G, Wase N, Shu S, Jenkins J, Zhou B, Torres-Rodríguez JV, Chen C, Sandor L, Plott C, Yoshinga Y, et al. 2022. Genome of Paspalum vaginatum and the role of trehalose mediated autophagy in increasing maize biomass. Nature communications 13: 7731.

74. Sun S, Zhou Y, Chen J, Shi J, Zhao H, Zhao H, Song W, Zhang M, Cui Y, Dong X, et al. 2018. Extensive intraspecific gene order and gene structural variations between Mo17 and other maize genomes. Nature genetics 50: 1289–1295.

75. Tesfamariam T, Yoshinaga H, Deshpande SP, Srinivasa Rao P, Sahrawat KL, Ando Y, Nakahara K, Hash CT, Subbarao GV. 2014. Biological nitrification inhibition in sorghum: the role of sorgoleone production. Plant and soil 379: 325–335.

76. Villegas D, Arevalo A, Nuñez J, Mazabel J, Subbarao G, Rao I, De Vega J, Arango J. 2020. Biological Nitrification Inhibition (BNI): Phenotyping of a Core Germplasm Collection of the Tropical Forage Grass Megathyrsus maximus Under Greenhouse Conditions. Frontiers in plant science 11: 820.

77. Wang X, Li Z, Policarpio L, Koffas MAG, Zhang H. 2020. De novo biosynthesis of complex natural product sakuranetin using modular co-culture engineering. Applied microbiology and biotechnology 104: 4849–4861.

78. Wang G, Zhang L, Guo Z, Shi D, Zhai H, Yao Y, Yang T, Xin S, Cui H, Li J, et al. 2023. Benefits of biological nitrification inhibition of Leymus chinensis under alkaline stress: the regulatory function of ammonium-N exceeds its nutritional function. Frontiers in plant science 14: 1145830.

79. Widhalm JR, Rhodes D. 2016. Biosynthesis and molecular actions of specialized 1,4- naphthoquinone natural products produced by horticultural plants. Horticulture research 3: 16046.

80. Zhang M, Gao X, Chen G, Afzal MR, Wei T, Zeng H, Subbarao GV, Wei Z, Zhu Y. 2023. Intercropping with BNI-sorghum benefits neighbouring maize productivity and mitigates soil nitrification and N2O emission. Agriculture, ecosystems & environment 352: 108510.

81. Zhang T, Huang W, Zhang L, Li D-Z, Qi J, Ma H. 2024. Phylogenomic profiles of whole- genome duplications in Poaceae and landscape of differential duplicate retention and losses among major Poaceae lineages. Nature communications 15: 3305.

