## Supplementary Figures for "Contrasting Rhizosphere Nitrogen Dynamics in Andropogoneae Grasses: Implications for Sustainable Agriculture"

**This PDF file includes:**

Supplementary Figure S1 to S7

**Other supplementary materials for this manuscript include the following:**

Supplementary Table S1 to S6

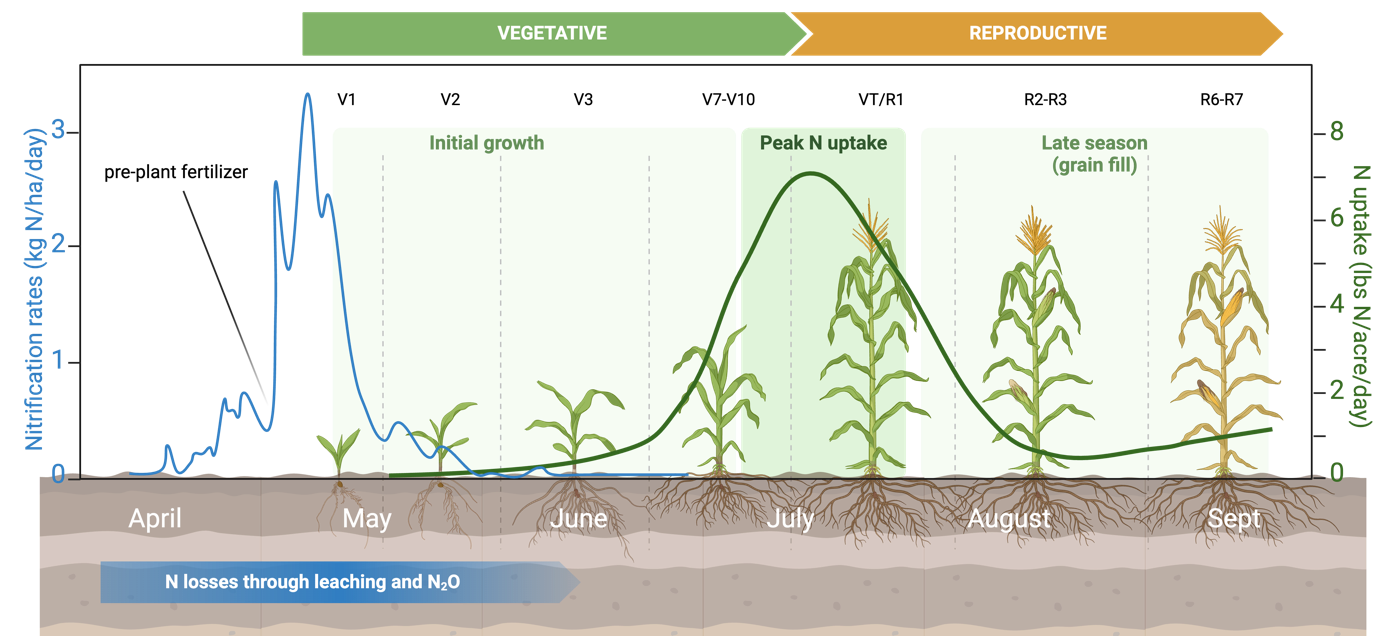

**Supplementary Figure S1. Mismatch in timing between crop fertilization, plant N demand and soil microbial activities.** The x-axis denotes maize growing season in Midwest US. Nitrification and nitrogen losses (y-axis on the left; blue) occurs before plant N uptake (y-axis on the right; green) can effectively compete for mineral N. Created with BioRender.com, after Bender et al (2013), Hartmann et al (2022), and Kosola et al (2023).

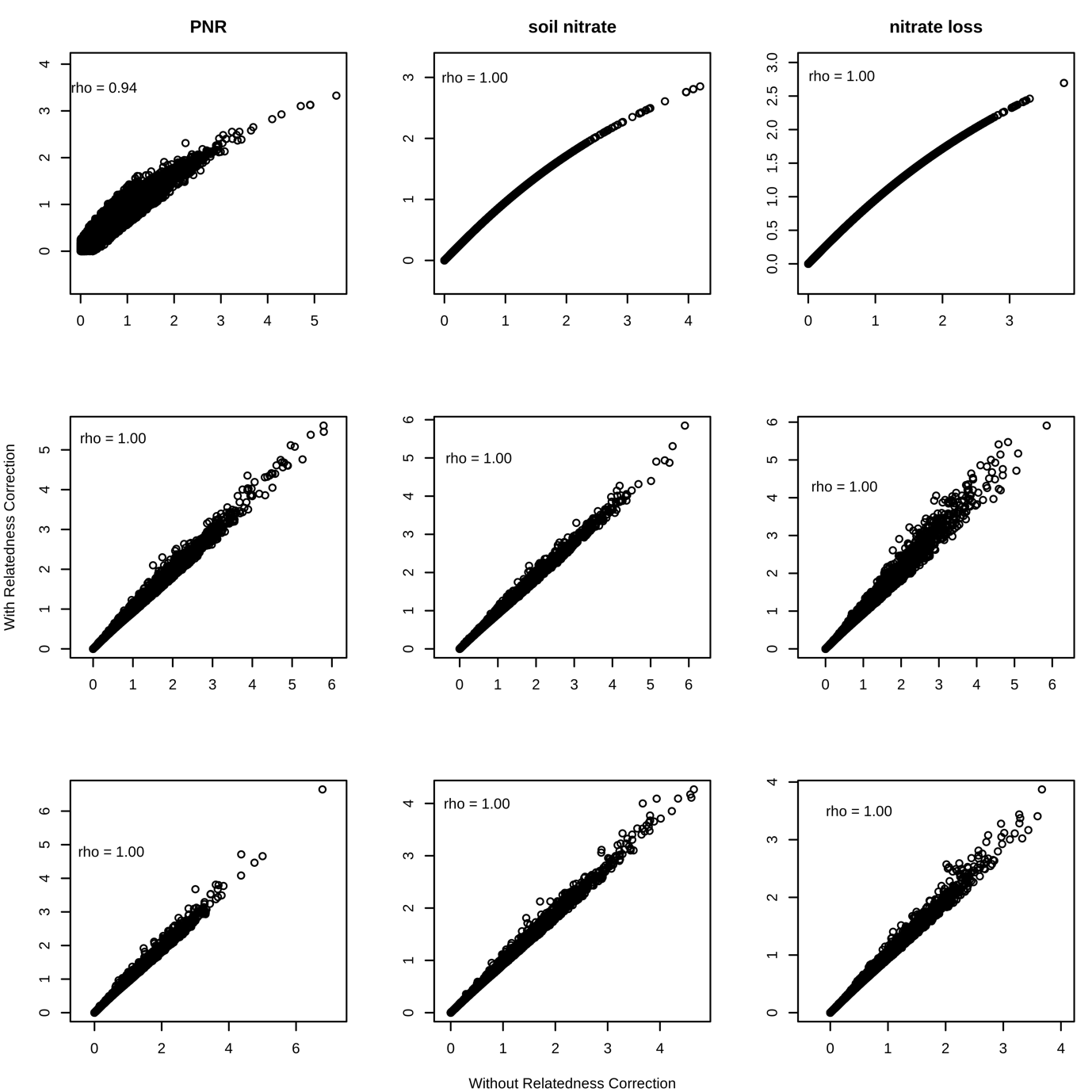

**Supplementary Figure S2. Significance of association test with and without accounting for taxa relatedness.** Significant correlation is detected between the significance (-log_10_ scale) with (y-axis) and without (x-axis) accounting for taxa relatedness in the model. Each column is a trait (PNR, soil NO_3_^-^ and NO_3_^-^ loss) and each row stands a homolog feature (LoF prediction, leaf expression and root expression)

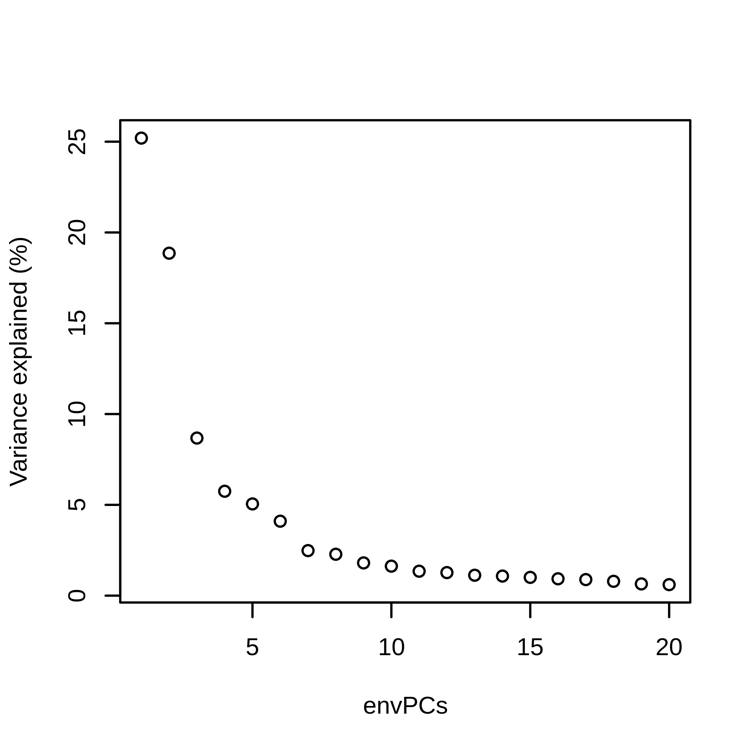

**Supplementary Figure S3. Variance explained by the top 20 envPCs.** In total, the top 20 envPCs explain >80% of total environmental variation.

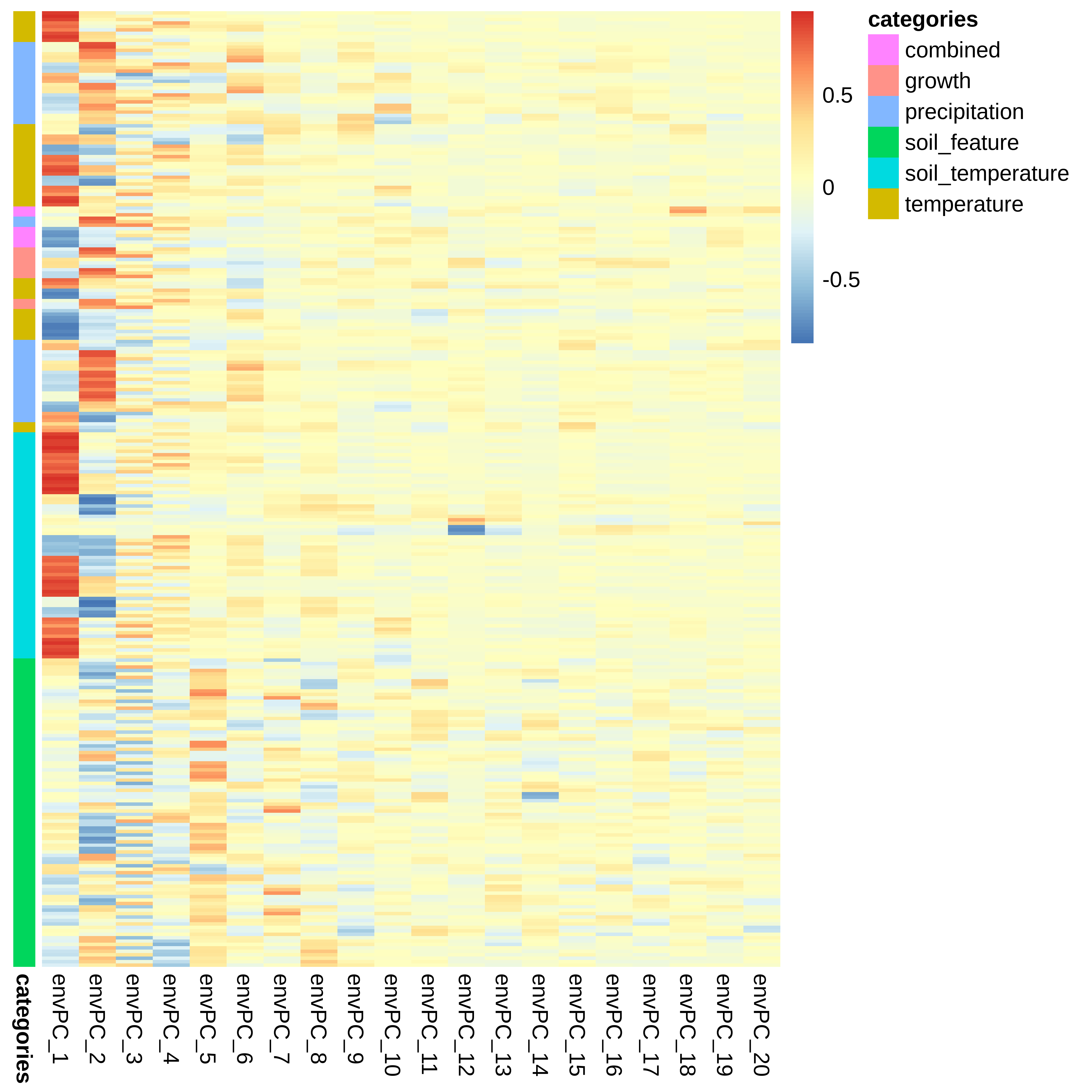

**Supplementary Figure S4. Correlation between each environmental feature and the top 20 envPCs.** Each row denotes one environmental features (classified into six categories). Each column denotes an envPC. The heat colors in each cell represent the correlation coefficients between each environmental feature and envPC. envPC_1 mainly correlates to temperature-related features; envPC_2 to precipitation-related features, the two major climatic factors. envPC_5 instead correlates to many soil features. envPC_12 correlates to specific soil temperature-related features (isothermality).

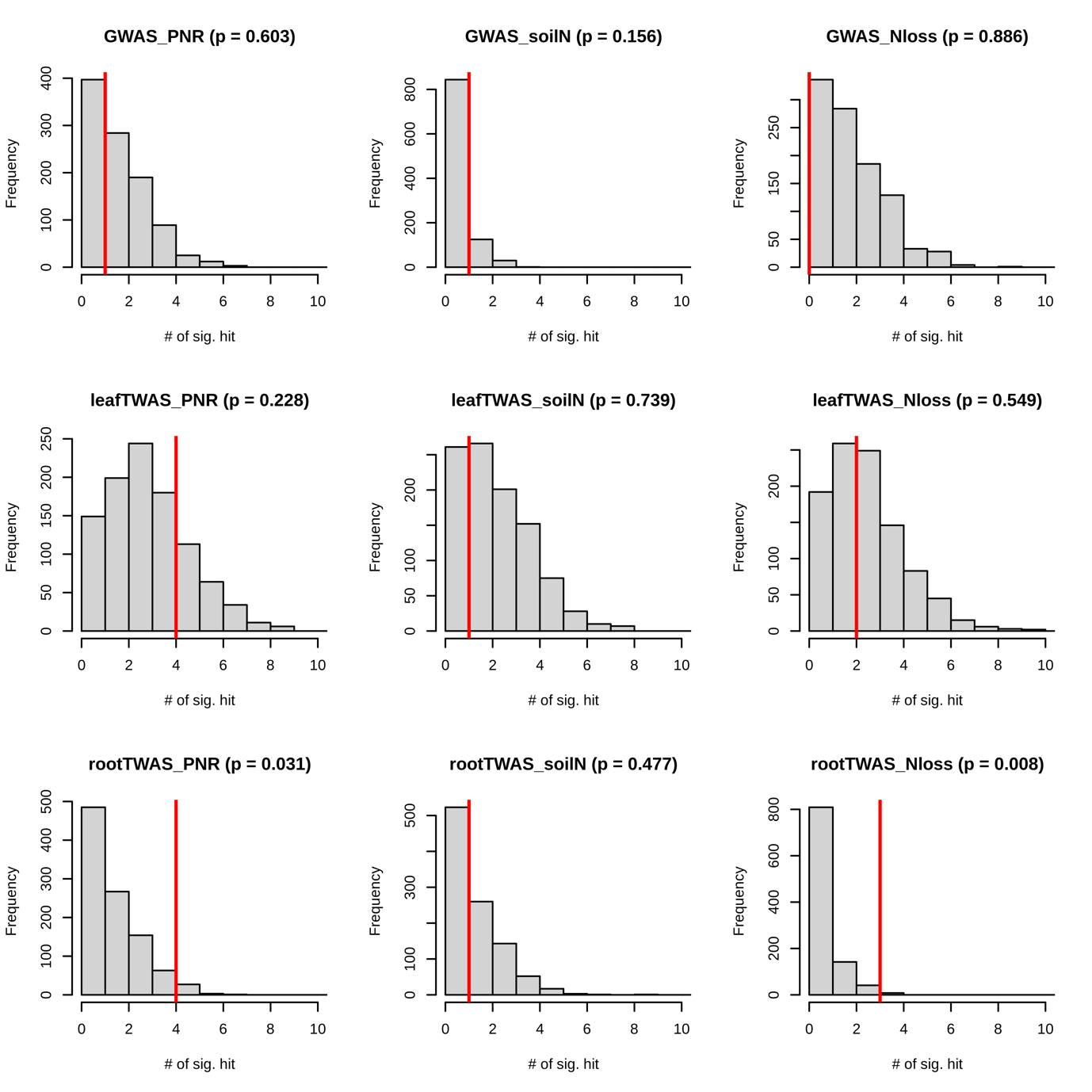

**Supplementary Figure S5. Enrichment of significant association signatures in a priori candidate gene set.** The grey distribution denotes null expectation of the number of significant association to detect in 1000 random sets of 188 homolog (X). The red vertical line indicates the empirical observation (Obs) and empirical p value is calculated as P(X>Obs). Each panel stands for the result of one association model between a trait and a homolog feature.

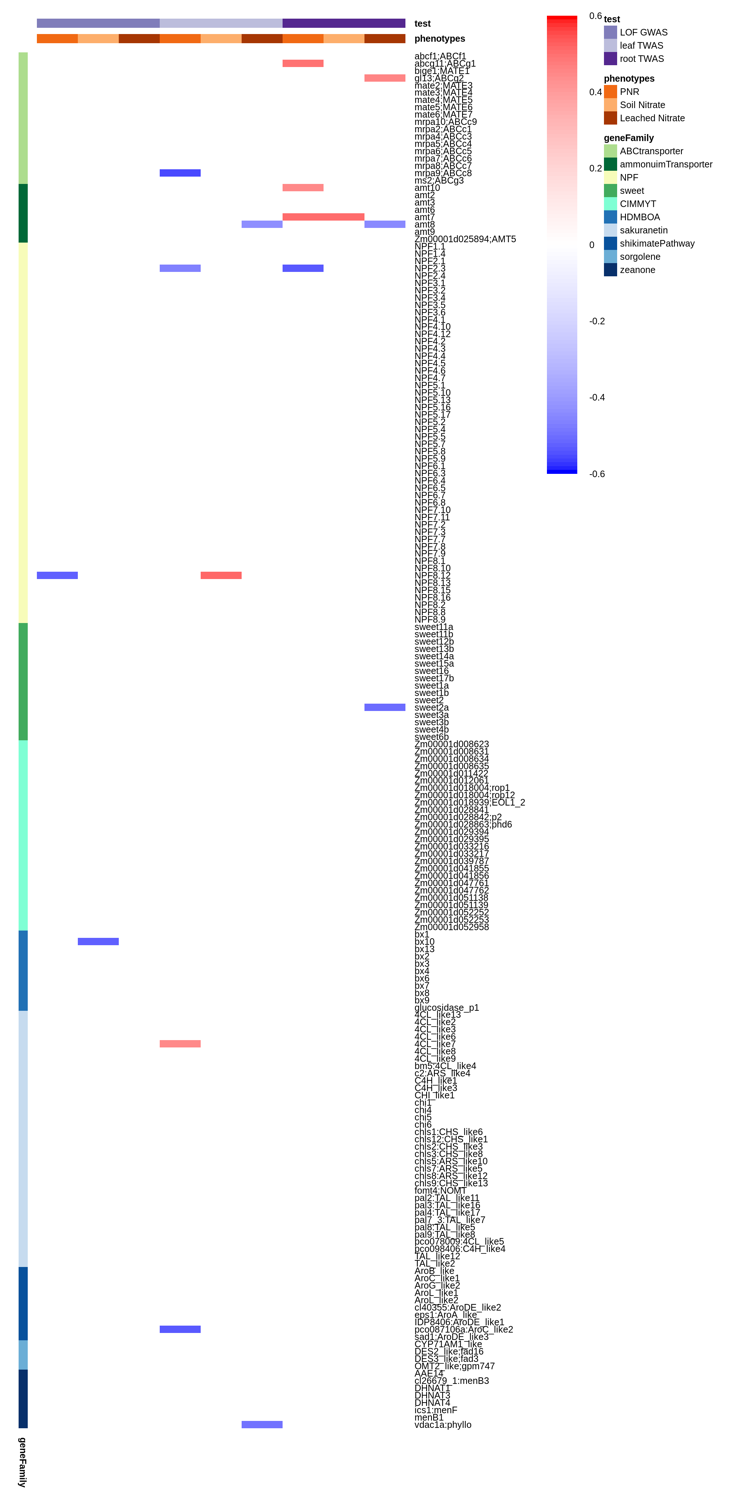

**Supplementary Figure S6. Summary of the cross-species association for the candidate homologs.** For each homolog, there are nine association tests (three genetic factors by three rhizosphere N traits) as denoted by the columns. Each row is a homolog that exhibits significant association in at least one trait-genetic combination. Row annotation colors indicate the gene classes each homolog belongs to. The cell colors denote the strength and direction of the association (Pearson’s correlation coefficient, r).

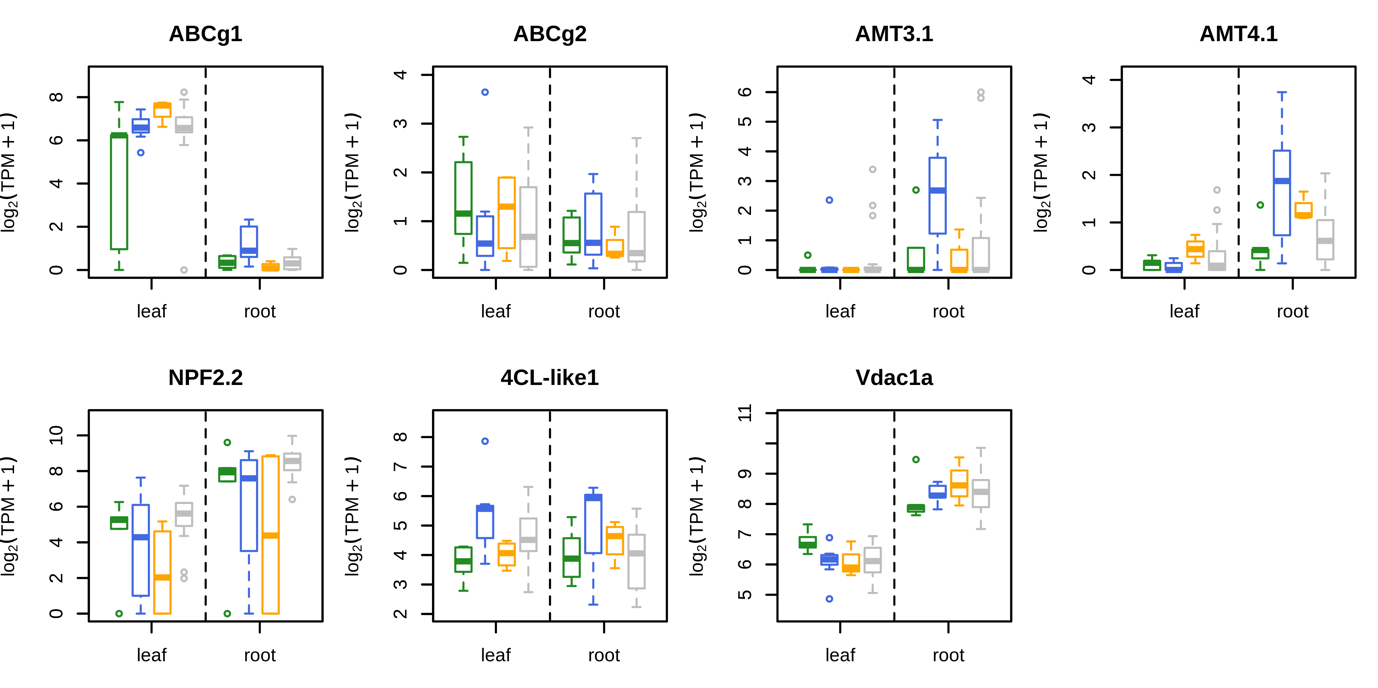

**Supplementary Figure S7. Homolog expression profiles of the significant candidates.** Candidate homolog expression in different tissues (left: leaf and right: root) in different species groups with distinct rhizosphere N traits (colors)
